# A short prokaryotic argonaute cooperates with membrane effector to confer antiviral defense

**DOI:** 10.1101/2021.12.09.471704

**Authors:** Zhifeng Zeng, Yu Chen, Rafael Pinilla-Redondo, Shiraz A. Shah, Fen Zhao, Chen Wang, Zeyu Hu, Changyi Zhang, Rachel J. Whitaker, Qunxin She, Wenyuan Han

## Abstract

Argonaute (Ago) proteins are widespread nucleic acid-guided enzymes that recognize targets through complementary base pairing. While in eukaryotes Agos are involved in RNA silencing, the functions of prokaryotic Agos (pAgos) remain largely unknown. In particular, a clade of truncated and catalytically inactive pAgos (short pAgos) lacks characterization. Here, we reveal that a short pAgo protein in *Sulfolobus islandicus*, together with its two genetically associated proteins, Aga1 and Aga2, provide robust antiviral protection via abortive infection. Aga2 is a membrane-associated toxic effector that binds anionic phospholipids via a basic pocket, which is essential for its cell killing ability. Ago and Aga1 form a stable complex that exhibits RNA-directed nucleic acid recognition ability and directly interacts with Aga2, pointing to an immune sensing mechanism. Together, our results highlight the cooperation between pAgos and their widespread associated proteins, suggesting an uncharted diversity of pAgo-derived immune systems that await to be discovered.

---

Argonaute (Ago) proteins are found across all domains of life and comprise a diverse family of defense elements against transposons, plasmids and viruses. Agos bind to short nucleic acid guides that direct the recognition of nucleic acid targets through complementary base-pairing (Hegge et al., 2018; Meister, 2013). Eukaryotic Agos (eAgos) use RNA guides to recognize RNA targets, which are then cleaved by either the eAgo’s intrinsic nuclease activity or through the recruitment of ancillary RNA-silencing factors (Hutvagner and Simard, 2008; Ketting, 2011). In contrast, many prokaryotic Ago (pAgo) proteins, including those from *Thermus thermophilus* (Swarts et al., 2014a), *Pyrococcus furiosus* (Swarts et al., 2015), *Methanocaldococcus jannaschii* (Zander et al., 2017) and *Clostridium butyricum* (Kuzmenko et al., 2020), perform DNA-guided target DNA interference. Previous works have shown that the DNA interference activity is essential for immunity against viruses and plasmids (Kuzmenko et al., 2020; Swarts et al., 2014a). In addition, a recent study reports that *T. thermophilus* Ago is involved in DNA replication completion, expanding the physiological roles of pAgos (Jolly et al., 2020). Some of the DNA-targeting pAgos also exhibit guide-independent DNase activity, which cleaves dsDNA and generates the DNA guides (Swarts et al., 2017; Zander et al., 2017). Comparatively, pAgos from *Marinitoga piezophila* (Kaya et al., 2016) and *Rhodobacter sphaeroides* (Miyoshi et al., 2016) use RNA guides to target DNA. Further, a pAgo from *Kurthia massiliensis* can employ both DNA and RNA guides to target DNA and RNA (Kropocheva et al., 2021; Liu et al., 2021c). Altogether, these studies highlight a remarkable diversity of guide and target preferences for pAgos.

Bioinformatic studies reveal that pAgos encompass substantially more diversity than eAgos (Makarova et al., 2009; Ryazansky et al., 2018). pAgos can be classified into three groups, i.e., long-A, long-B and short. Long-A and long-B pAgo proteins share similar domain architecture with eAgos, containing six structural segments: N-terminal, L1 (Linker 1), PAZ (PIWI-Argonaute-Zwille), L2 (linker 2), MID (Middle) and PIWI (P-element Induced Wimpy Testis) domains. The PIWI domain contains a characteristic RNaseH fold and comprises the nuclease domain. The PIWI domain of long-A pAgos is functionally active, endowing them target cleavage ability and most characterized pAgos belong to this group. Due to their nuclease activity, long-A pAgos have been repurposed for nucleic acid detection (He et al., 2019; Liu et al., 2021b). In contrast, long-B pAgos, represented by *Rhodobacter sphaeroides* (Rs) Ago, appear to harbor mutations in the cognate catalytic residues that render them inactive nucleases (Kaya et al., 2016). Short pAgo proteins only contain the MID and PIWI domains and, analogous to the long-B group, are inactive nucleases due to mutations within PIWI domain. Currently, the functions and molecular mechanisms driving short pAgos remain unknown.

Interestingly, previous computational analyses have identified many gene families frequently encoded in the genomic neighborhoods of pAgos and diverse protein domains directly fused with pAgos (Makarova et al., 2009; Ryazansky et al., 2018; Swarts et al., 2014b). Although the associated proteins are predicted to be functionally linked with pAgos, little is known about their physiological roles and mechanisms of action. In this study, we report that a short pAgo from *Sulfolobus islandicus* and its associated proteins collaborate to provide robust antiviral immunity by mediating an abortive infection response.

## Results

### SiAgo system provides anti-viral defense via abortive infection

*S. islandicus* M164 and other related strains (Reno et al., 2009) encode a pAgo protein (M164_1614), which is non-essential for cell viability (Zhang et al., 2018). Analysis of the genetic neighborhood of *M164_1614* revealed genes commonly associated with mobile genetic elements (Figure S1A), suggesting that the region is mobile or an integration hotspot for horizontally acquired elements. In support of this, the entire region is absent in closely related strains, e.g., in *S. islandicus* M1425 (Figure S1B). Notably, although the genes in this region vary across *S. islandicus* strains, *ago* is invariably associated with three other genes i.e., *M164_1612, M164_1613* and *M164_1615* (Figure 1A, Figure S1A), suggestive of a functional connection. We refer to these syntenic genes as SiAgo system hereafter.

**Figure 1.**
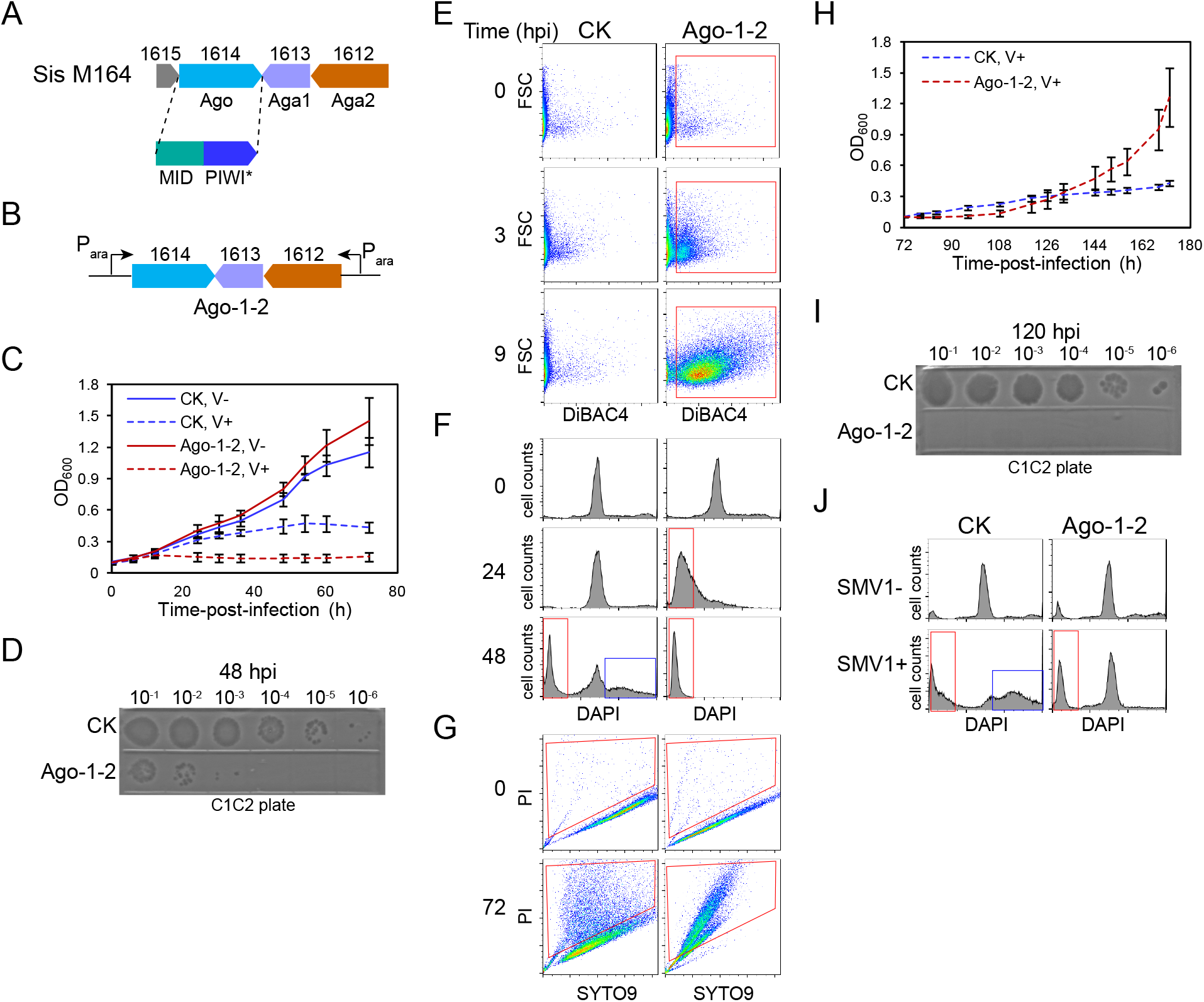
SiAgo system defends against SMV1 by mediating Abi. (A) Schematic of the Ago system from *S. islandicus* M164. The gene locus tags are indicated above the cluster. M164_1614 encodes a short Ago protein containing a MID domain and a catalytically inactive PIWI domain (PIWI*). (B) Schematic of the inducible cassettes reconstituting the SiAgo system in the pSeSD-Ago-1-2 expression plasmid. (C) Growth curves of the *S. islandicus* strains containing the SiAgo system (Ago-1-2), or lacking it (CK), during the course of SMV1 infection at a MOI of ∼2 (V+) in the sucrose medium. Isogenic cultures without SMV1 virus (V-) are shown as controls. The data show means of three independent replicates. Error bars indicate the standard deviations. (D) Viral titers in the cultures post infection. The supernatant of the cultures was serially diluted and spotted onto plates carrying *S. islandicus* C1C2 cells at 48 hpi. (E) Analysis of the cell membrane polarity by DiBAC_4_(3) staining. The cells were stained by DiBAC_4_(3) and analyzed by flow cytometry at indicated time points post SMV1 infection. The data are shown in the green fluorescence (DiBAC_4_(3) signal, horizontal axis)-FSC (forward scattered light, vertical axis) cytograms. DiBAC_4_(3)-positive cells are indicated by red boxes. (F) Analysis of the DNA content distributions by DAPI staining. The cells were stained by DAPI and analyzed by flow cytometry at indicated time points post viral infection. The data are shown in the blue fluorescence (DAPI signal, horizontal axis)-cell count (vertical axis) histograms. Cells exhibiting low DNA content are indicated in red boxes, while cells containing DNA content >2 chromosomes are indicated in blue boxes. (G) Analysis of the cell membrane permeability by SYTO9/PI staining. The cells were stained by SYTO9 and PI simultaneously and analyzed by flow cytometry at 72 hpi. The data are shown in the green fluorescence (SYTO9 signal, horizontal axis)-red fluorescence (PI signal, vertical axis) cytograms. PI-positive cells are indicated in red boxes (H) Growth curves of the infected cultures which were transferred into fresh medium at 72 hours post infection (hpi). Means of three independent replicates are shown. Error bars indicate the standard deviations. (I) Viral titers in the infected cultures at 120 hpi. (J) DNA content distributions of the infected (SMV1+) and uninfected (SMV1-) cultures at 120 hpi.

Protein sequence alignment of SiAgo with previously characterized long-A pAgos revealed that SiAgo only contains the MID and PIWI domains (Figure 1A, Figure S1C) and thus belongs to the short pAgo family. Notably, the PIWI domain of SiAgo is mutated, indicating that it lacks nucleic acid target cleavage capabilities (Figure S1C). HHpred analysis (Soding et al., 2005) of the associated proteins suggests *M164_1615* might play a regulatory role as it contains a HTH DNA binding domain with high similarity to diverse transcription factors (Figure S1D), while M164_1612 and M164_1613 yielded no significant hits to protein domains of known function. In this study, we aimed to investigate the putative coordinated functions of SiAgo, M164_1613 (SiAgo-associated protein1, SiAga1) and M164_1612 (SiAga2).

For this purpose, we expressed SiAgo, SiAga1 and SiAga2 using the pSeSD shuttle vector in *S. islandicus* E233S1 (Figure 1B), a genetic host derived from *S. islandicus* Rey15A that does not encode pAgo homologs (Deng et al., 2009). In pSeSD, expression of the genes of interest is driven by an engineered arabinose promoter (P_ara_) that allows tunable protein expression (Peng et al., 2012). We then infected the strain expressing the SiAgo system (Ago-1-2) and a control strain carrying the empty pSeSD plasmid (CK) with the *Sulfolobus* SMV1 virus that can infect and replicate in *S. islandicus* E233S1 (Guo et al., 2019; Uldahl et al., 2016), and assessed culture growth dynamics and viral proliferation post infection (Figure 1C and D). In the absence of SMV1, expression of Ago-1-2 did not exert an observable effect on culture growth. Upon SMV1 infection at an MOI (multiplicity of infection) of ∼2, the initial growth kinetics of CK were substantially retarded, yet the OD_600_ (optical density at 600 nm) slowly increased up to ∼0.4. In contrast, the growth of the Ago-1-2 strain was almost completely inhibited (Figure 1C). Measurements of the viral titers in the cultures with plaque forming unit (PFU) assay using a sensitive strain (*S. islandicus* C1C2)(Uldahl et al., 2016) indicates that the virus successfully replicated in the CK culture but not in the strain containing the SiAgo system (Figure 1D). Altogether, the data suggest that expression of Ago and its associated proteins inhibits virus proliferation by mediating cell death or dormancy.

To gain further insights into the effects of the SiAgo system on SMV1-infected cells, we stained the cell populations with three dyes (DiBAC_4_(3), DAPI and a mixture of SYTO9 and PI) at different time points post infection. While DiBAC_4_(3) is an indicator of the cell membrane depolarization (Bortner et al., 2001; Erental et al., 2012), DAPI stains DNA and indicates cellular DNA content distributions. In contrast, SYTO9 is able to enter live cells with an intact membrane and PI only enters dead cells that have lost membrane integrity (Leuko et al., 2004; Bize et al., 2009). Analysis of the stained cells with flow cytometry indicated that SMV1-infected Ago-1-2 cells started to lose membrane polarity at 3 hpi and that most cells lost membrane polarity at 9 hpi, while SMV1 infection did not induce apparent cell membrane depolarization in CK culture at 9 hpi (Figure 1E). Furthermore, infected Ago-1-2 cells were depleted of DNA at 24 hpi, whereas the SMV1-infected CK cells showed a similar DNA content distribution to uninfected cells (Figure 1F). At 48 hpi, a fraction of the infected CK cells had lost their genomic DNA, while a larger fraction contained more DNA than uninfected cells, suggesting an accumulation of viral DNA and/or suppression of cell division (Guo et al., 2019; Liu et al., 2021a). Finally, SYTO9/PI staining indicated that most infected Ago-1-2 cells lost membrane integrity at 72 hpi, in contrast to the much smaller fraction of infected CK cells and in line with the DNA content distribution analysis (Figure 1G). Together, these results indicate that SiAgo system kills infected cells to inhibit viral proliferation.

The defense strategy that kills infected cells so that the uninfected clonal cells can grow in a virus-free environment is referred to as abortive infection (Abi) (Isaev et al., 2021; Lopatina et al., 2020). To further investigate whether the SiAgo system confers such a fitness advantage via an Abi response, we transferred the infected cultures into fresh medium at an OD_600_ of ∼0.1 at 72 hpi and continued to monitor their growth until 172 hpi. Indeed, the growth of the strain carrying the SiAgo system was restored after 108 hpi (Figure 1H) and less viral particles were observed in comparison to before the transfer (Figure 1I). In agreement with the clearance of infection at this time point, flow cytometry analysis revealed that a considerable fraction of cells showed similar DNA content distributions to uninfected cells (Figure 1J). The data confirm that SiAgo system allows the cell culture to successfully clear the viral infection and restore a population of virus-free cells. In contrast, the CK culture displayed long-lasting slow growth kinetics and continued to exhibit high-titer virus particles. The DNA content distributions of the CK culture were also similar to those before the transfer, indicating that SMV1 maintains a stable infectious status in the cells instead of inducing cell lysis, in line with previous studies on SMV1 and other *Sulfolobus* viruses (Guo et al., 2019; Liu et al., 2021a; Prangishvili et al., 2006; Uldahl et al., 2016).

### Defense requires all three proteins and SiAga2 is the toxic effector

To investigate how the SiAgo system performs Abi, we firstly analyzed the minimum components required for the process. We constructed strains expressing combinations of only two of the three proteins, i.e., Ago-Aga1, Ago-Aga2 and Aga1-Aga2. The corresponding strains, as well as the strains containing empty vector (CK) and expressing all three components (Ago-1-2), were subjected to SMV1 infection and subsequent phenotypic analysis. Importantly, the results show that lack of any one of the three components abolished DNA depletion in cells and resulted in successful viral proliferation (Figure 2A and B), highlighting that all three proteins are essential for the execution of the Abi response.

**Figure 2.**
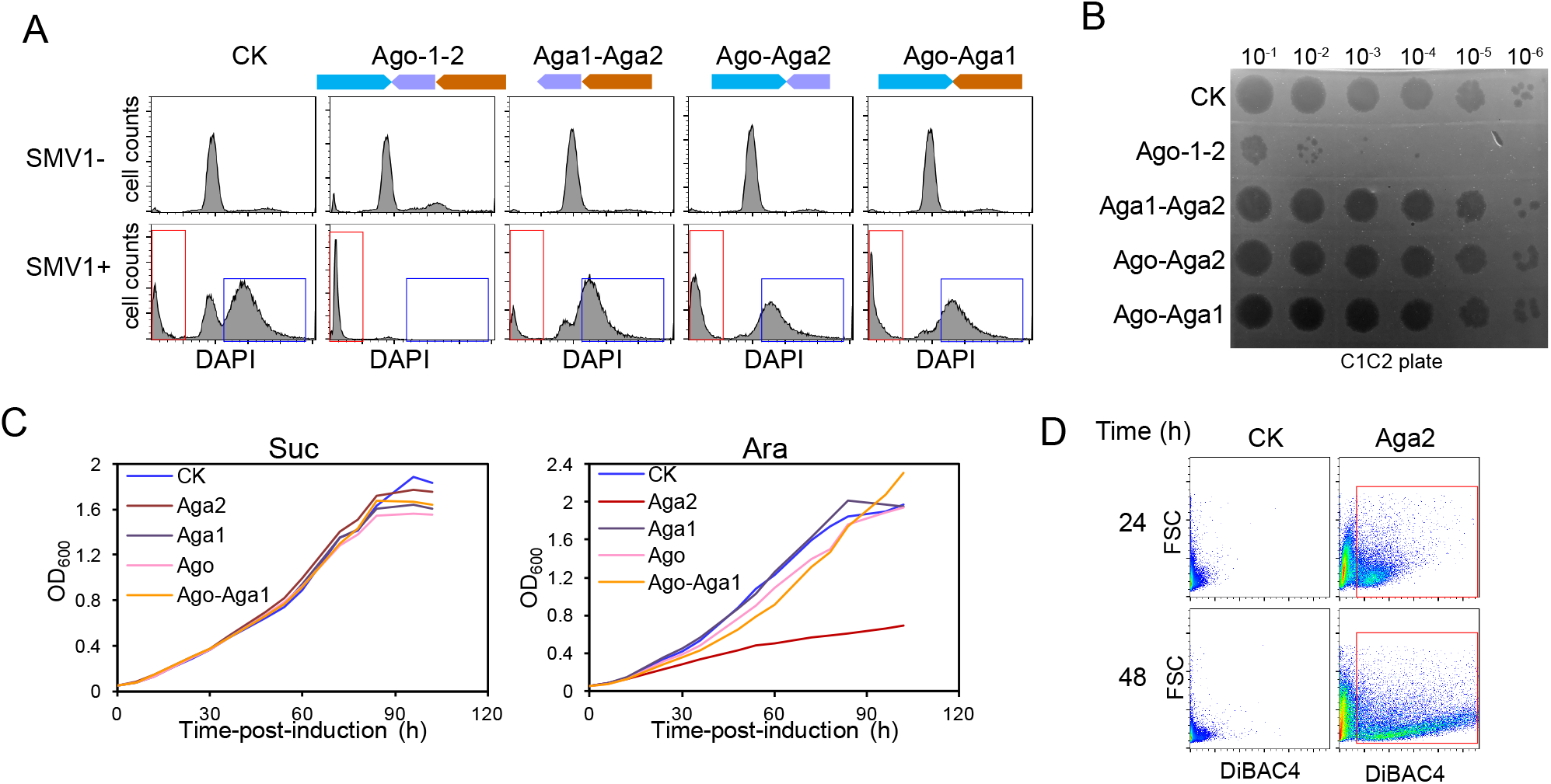
SiAgo, SiAga1 and SiAga2 are required for Abi execution and SiAga2 acts as toxic effector. (A) Strains containing the complete SiAgo system or incomplete versions of the system (each lacking one of the three proteins), or lacking the system (CK) were grown in the presence or absence of SMV1 (SMV1+ and SMV1-, respectively). DNA content distributions were analyzed at 48 hpi. DNA-less cells are indicated in red boxes, while cells containing DNA content >2 chromosomes are indicated in blue boxes. (B) Viral titers of the cultures derived from the strain versions depicted in (A) at 48 hpi. (C) Growth curves of the strains expressing different components of the SiAgo system in sucrose (Suc) and arabinose (Ara) medium, respectively. The strains include CK and those expressing Aga1, Aga2, Ago, Ago-Aga1, respectively. The data of one out of two independent replicates are shown. (D) Analysis of the cell membrane polarity of the cells overexpressing Aga2 or not (CK)

In order to mediate anti-viral protection, Abi defense systems must include a sensor module and a killer module (Lopatina et al., 2020); the former senses viral infection and activates the latter to induce cell toxicity. First, we sought to reveal which of the three protein components is the toxic effector. To address this question, we analyzed whether overexpression of any of the proteins would induce cell toxicity. Each of the three proteins was expressed using the P_ara_ promoter and the expression strains were cultured in media containing either sucrose (Suc) or arabinose (Ara), the latter of which can greatly induce the expression of the proteins of interest (Peng et al., 2012). In the Suc medium, the growth of the three strains was similar to the CK strain (Figure 2C, Figure S2A). However, while in the Ara medium the strains expressing SiAgo or SiAga1 showed normal growth kinetics, overexpression of SiAga2 resulted in significant growth retardation, indicative of its toxicity (Figure 2C, Figure S2A). Moreover, DiBAC_4_(3) staining revealed that overexpressed SiAga2 induced membrane depolarization (Figure 2D). On the other hand, even co-overexpression of SiAgo and SiAga1 did not exhibit any inhibition on culture growth, further reinforcing the notion that SiAga2 is the effector of SiAgo system and can perform cell toxicity independently.

### SiAga2 binds to anionic head groups of phospholipids

Next, we sought to explore the molecular mechanisms underlying the SiAga2-mediated Abi antiviral immune response. Analysis of the protein sequence using TMHMM server (Krogh et al., 2001) indicated that SiAga2 contains two transmembrane regions (2×TM) (Figure 3A). We subsequently confirmed that SiAga2 is a membrane-associated protein with fluorescence *in situ* hybridization *in vivo* (Figure 3B). Then, we predicted the protein structure of SiAga2 leveraging the AlphaFold2 structure prediction pipeline (Jumper et al., 2021) (Figure 3C). The results indicate that the soluble domain of SiAga2 contains a basic pocket facing the membrane. We hypothesized that such a domain architecture could be responsible for mediating the interaction with the anionic head groups of membrane lipids (Cho and Stahelin, 2005). To this end, we expressed SiAga2ΔC (lacking 2×TM) and screened for potential targets using commercial strips carrying a panel of immobilized lipids. Noticeably, the membrane lipids of archaea are significantly different from their bacterial and eukaryotic counterparts, particularly in the carbon chains, the bound linking carbon chains and glycerol, and the position of phosphate group on glycerol (Lombard et al., 2012). However, the head groups of membrane lipids are shared across the three domains of life. Specifically, the head groups of *Sulfolobus* include inositol, ethanolamine and glycerol (Koga and Morii, 2005), all of which are included in the tested lipids.

**Figure 3.**
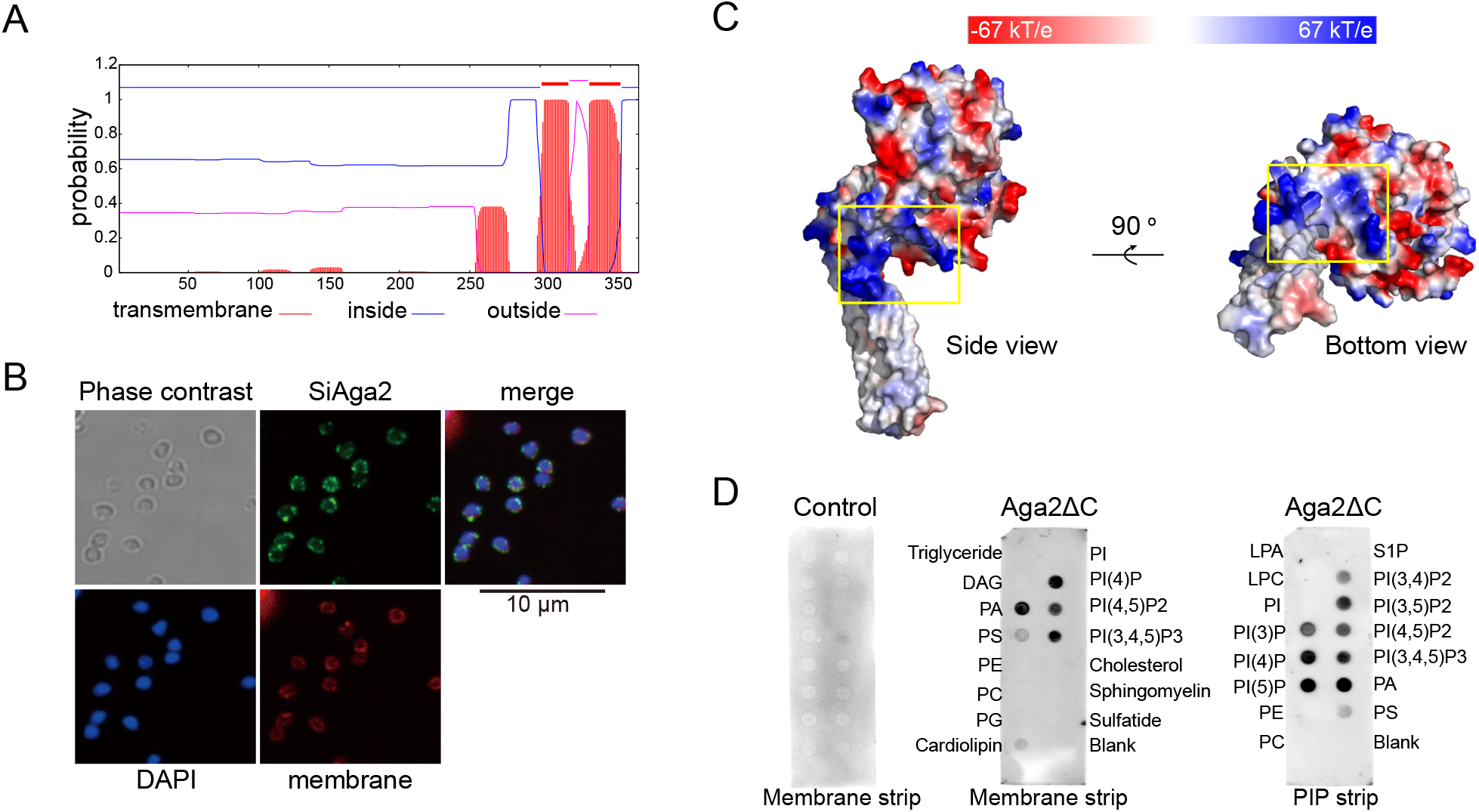
SiAga2 is a membrane protein with affinity to anionic phospholipids. (A) Prediction of transmembrane topology of SiAga2 using TMHMM Server. (B) Fluorescence microscopy analysis of subcellular localization of SiAga2. Images show phase contrast, DAPI staining of DNA (blue), SiAga2 (green), membrane (red) and merged images. (C) Surface representation of the structural model of SiAga2 colored according to electrostatic surface potential. The predicted basic pocket is indicated in yellow boxes. Left: side view; right: bottom view. (D) Binding of SiAga2ΔC to the membrane and PIP strips. The strips were incubated with SiAga2ΔC or not (control), and the bound SiAga2ΔC was detected by western blot.

The strip binding screening revealed that SiAga2ΔC strongly bound to phosphatidic acid (PA) and the lipids containing phosphorylated phosphoinositide (phosphatidylinositol (n)-phosphate, PI(n)P), and exhibited lower affinity to cardiolipin and phosphatidylserine (PS) (Figure 3D). Comparison of the affinitive and non-affinitive lipids indicated two apparent rules dictating substrate preference for SiAga2ΔC (Figure S3). First, the phosphate group is important, as indicated by the comparison of the PI(n)P and phosphatidylinositol (PI) (Figure S3A), and the comparison of PA and diacylglycerol (DAG) (Figure S3B). Phosphatidylcholine (PC), phosphatidylethanolamine (PE), and phosphatidylglycerol (PG) are not bound probably because their phosphate group is covered by other moieties. Second, SiAga2ΔC binds to phospholipids instead of lysophospholipids, since sphingosine 1-phosphate (S1P) and lysophosphatidic acid (LPA) (which contain a free phosphate group), are non-affinitive lipids (Figure S3C).

### The basic pocket of SiAga2 is essential for mediating Abi and binding anionic phospholipids

We further explored the functional implications of the identified basic pocket. To analyze whether the pocket is important for mediating Abi, we constructed strains expressing versions of the SiAgo system where SiAga2 contains substitution mutations of the alkaline residues predicted to be within or close to the pocket, i.e., R7A-R8A (M1), K12A-K13A (M2) and R7A-R8A-K12A-K13A (M3) (Figure 4A). Analysis of the response of the strains to SMV1 infection indicates that mutation of R7-R8 and K12-K13 significantly impaired membrane depolarization, while mutation of the four residues (tetramutation) abolished the phenomenon entirely (Figure 4B). In support of this, the DNA content distributions of the tetramutation strain were similar to that of the CK strain lacking the SiAgo system, indicating that the tetramutation abolished DNA loss (Figure 4C). Moreover, the tetramutation also lost the inhibition of viral proliferation, while each dimutation moderately impaired the inhibition of viral proliferation (Figure 4D). Together, the data indicate that the basic pocket in SiAga2 is essential for mediating Abi. To further analyze whether the mutations would affect the affinity of SiAga2ΔC to lipids, we expressed the corresponding SiAga2ΔC mutants and analyzed their affinity to the membrane strip. The results showed that each dimutation largely impaired the affinity to PA and PI(n)Ps, and that the tetramutation almost abolished the lipid binding ability (Figure 4E), indicating that the basic pocket is indeed involved in binding to the anionic head groups of phospholipids.

**Figure 4.**
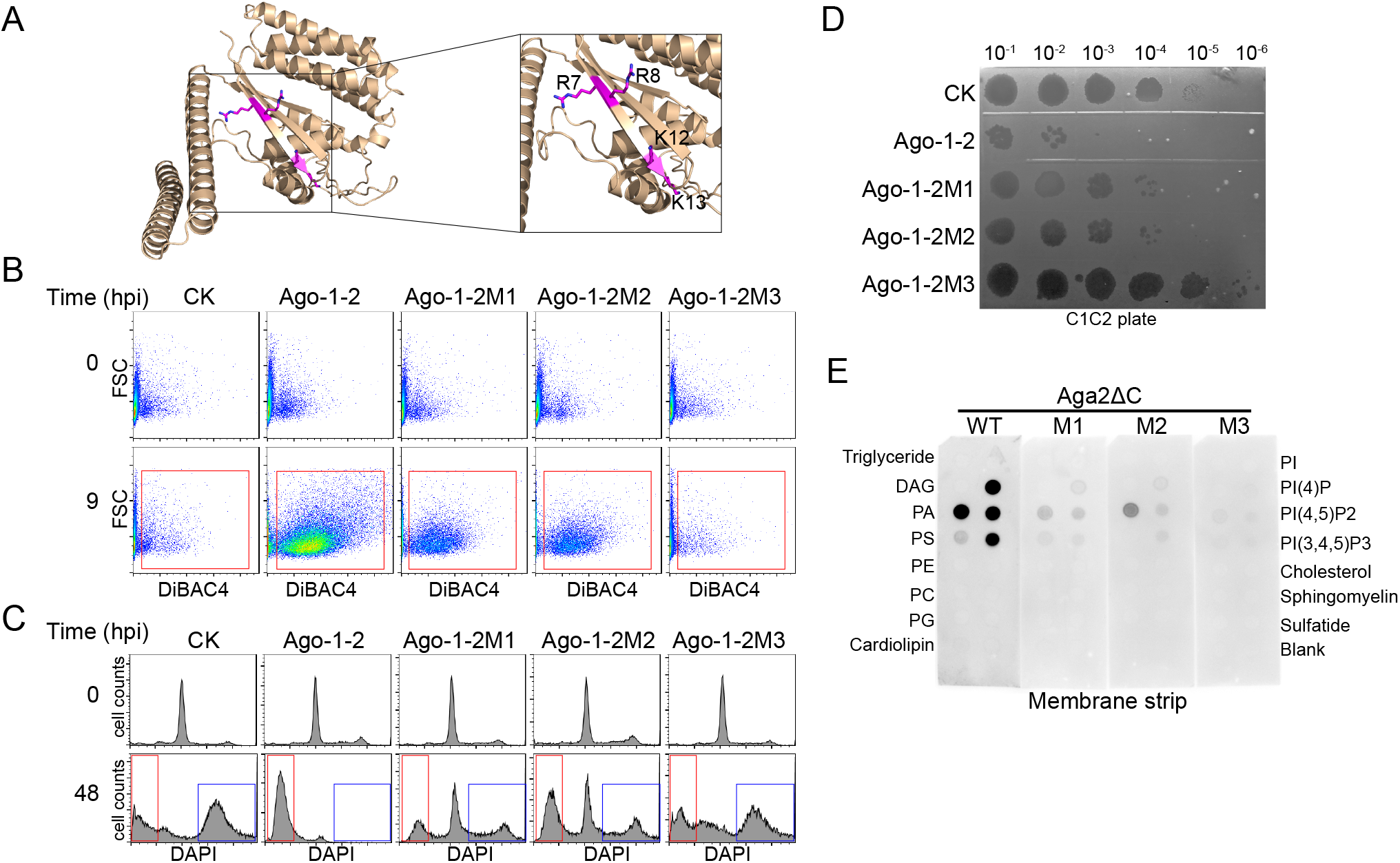
The basic pocket of SiAga2 is crucial for the Abi response and binding to anionic phospholipids. (A) The overall view and a close-up view of the basic pocket of SiAga2. The indicated residues were selected for mutagenesis. The protein is displayed in the same orientation as in Figure 3C bottom view. (B) Membrane depolarization of the cultures containing wild type (WT) or mutated SiAgo systems at 0 and 9 h post SMV1 infection. The mutations are within SiAga2: R7A-R8A (M1), K12A-K13A (M2) and R7A-R8A-K12A-K13A (M3). CK denotes the control lacking the SiAgo system. (C) DNA content distributions of the cultures containing WT or mutated SiAgo systems at 48 h post SMV1 infection. (D) Viral titers of the cultures containing WT or mutated SiAgo systems at 48 h post SMV1 infection. (F) Binding of SiAga2ΔC (WT) and its mutants to the membrane strip.

### SiAgo and SiAga1 interact with SiAga2 and enhance its toxicity

Given that SiAga2 is the sole toxic protein, we propose that SiAgo and SiAga1 are likely implicated in the sensor module in the Abi process. To gain insights into the functions of these two proteins, we compared the phenotypes of the strains expressing Aga2 or co-expressing Ago-1-2 in the Ara medium. The results show that, although overexpression of SiAgo and SiAga1 exerted little effects on culture growth or membrane polarity, co-overexpression of Ago-1-2 resulted in slower culture growth and more membrane-depolarized cells than overexpression of SiAga2 alone (Figure 5A and B, Figure S2B). The data therefore indicate that overexpressed SiAgo and SiAga1 enhance the toxicity of SiAga2 in the absence of viral infection.

**Figure 5.**
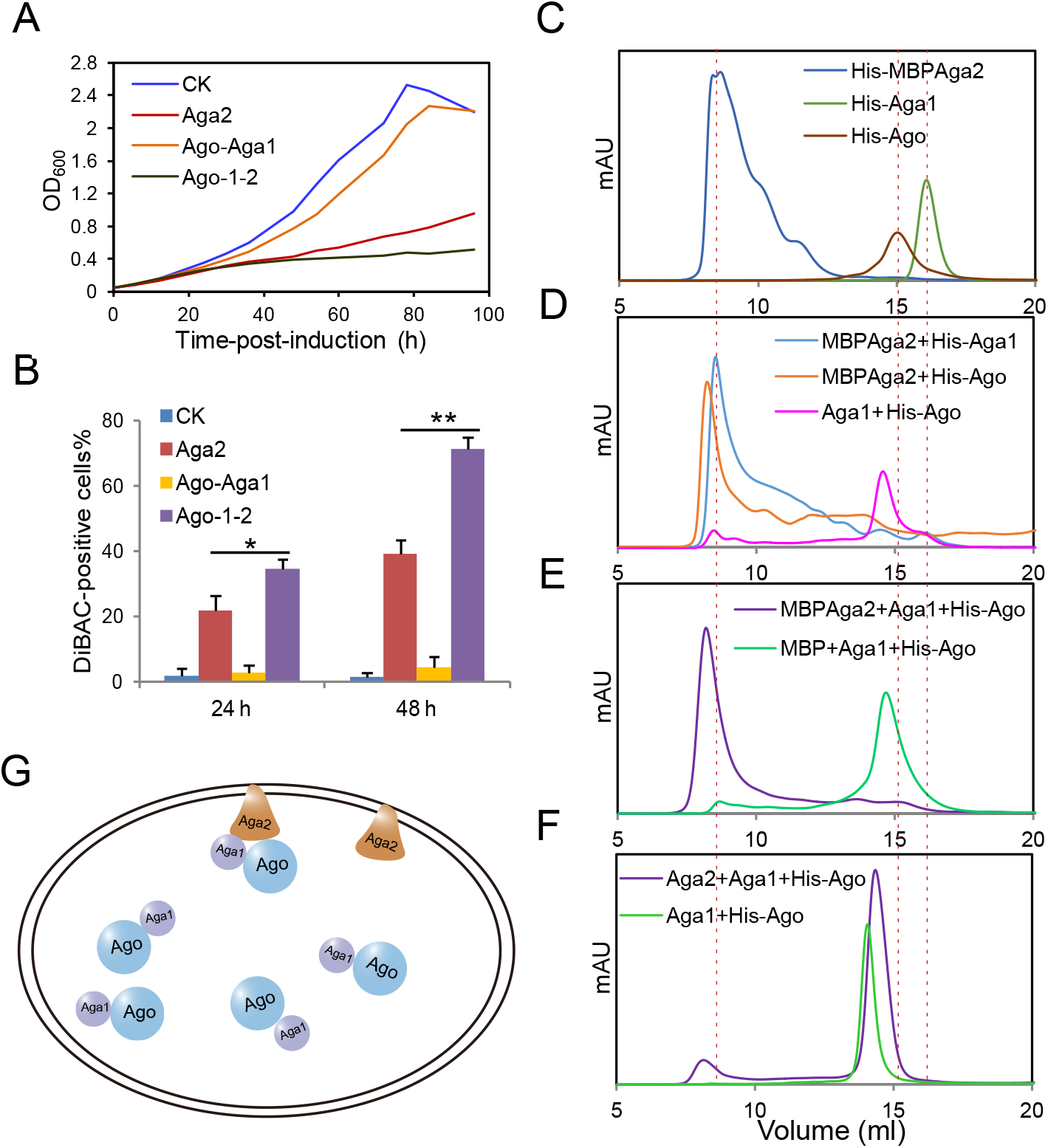
Interplay between SiAgo, SiAga1 and SiAga2. (A) Growth curves of the strains lacking the SiAgo system (CK), expressing Aga2, Ago-Aga1 or all the three proteins (Ago-1-2) in Ara medium. The data of one of three independent replicates are shown. (B) Quantification of the membrane-depolarized cells in the strains. Data shown are means of three independent repeats ± standard deviation. *: p<0.05; **: p<0.01. (C) Gel filtration analysis of purified His-MBPAga2, His-Aga1 and His-Ago, respectively. (D) Gel filtration analysis of co-expressed MBPAga2+His-Aga1, MBPAga2+His-Ago, and Aga1+His-Ago, respectively. (E) Gel filtration analysis of co-expressed MBPAga2+Aga1+His-Ago and MBP+Aga1+His-Ago, respectively. (F) Gel filtration analysis of co-expressed Aga2+Aga1+His-Ago and Aga1+His-Ago from *Sulfolobus* cells, respectively. (G) A model depicting the interactions between the three proteins in *Sulfolobus* cells.

Signal transduction from sensing viral infection to the activation of the toxic effector requires a specific interplay between the sensor and killer modules. To reveal possible interactions between SiAgo, SiAga1 and SiAga2, the three proteins were co-expressed together or in pairs in *E. coli*, or expressed independently as controls. In the co-expression strains, only one protein was labeled with a 6×His tag and used as the bait during purification in order to check whether the other proteins were co-eluted. Full length SiAga2 was expressed as a fusion with the maltose binding domain (MBP) to increase its stability. The purification procedure included Ni-NTA chromatography, anion exchange chromatography and size exclusion chromatography (SEC). During SEC, the elution volume of SiAga1 and SiAgo was about 16.1 and 15.0 mL respectively. MBP-fused SiAga2 was eluted at ∼8.6 mL even though the sample has been treated with detergent, indicative of a large protein aggregate (Figure 5C, Figure S4A∼C). Purification of co-expressed His-Ago+Aga1 resulted in a single peak containing both proteins at about 14.5 mL (Figure 5D, Figure S4D), indicative of a stable complex. Estimation of the stoichiometry of the two proteins suggests a 1:1 ratio in the complex (Figure S4E and F). In addition, MBPAga2 was co-purified with His-Ago and His-Aga2, respectively, resulting in that a fraction of His-Ago and His-Aga2 was eluted at around 8.5 mL (Figure 5D, Figure S4G and H). Similarly, purification of co-expressed His-Ago+Aga1+MBPAga2 yielded a single peak at ∼8.2 mL, which contains all the three proteins (Figure 5E, Figure S4I). As negative control, purification of co-expressed His-Ago+Aga1+MBP did not obtain MBP (Figure 5E, Figure S4J), excluding the possibility that His-Ago+Aga1 interacts with MBP. Together, the data indicate that the three components interact with each other in pairs and can form a ternary complex.

To reveal how they interact with each other in the native host, we expressed and purified the three proteins from *Sulfolobus* cells using His-tagged SiAgo as the bait. SEC results indicate that the co-purification resulted in a peak at ∼14.5 ml, representing the His-Ago+Aga1 complex and a small peak at ∼8.2 mL containing all the three proteins (Figure 5F, Figure S4K∼N). The data reinforce the conclusion that the three proteins form a ternary complex and support that SiAgo and SiAga1 form a complex in the cytoplasm, which are then recruited to SiAga2 in the native host (Figure 5G).

### Guide and target binding of Ago-Aga1 complex

Ago proteins are inherently directed by nucleic acid guides to bind and/or cleave complementary nucleic acid targets. We thus predicted that the nucleic acid recognition ability is important for the function of the SiAgo system. To shed light on how SiAgo binds to nucleic acids, we first performed an electrophoretic mobility shift assay (EMSA) using single strand RNA and DNA substrates. Because the 5’ group of the guide strand can affect the recognition by Ago proteins (Kaya et al., 2016; Ma et al., 2005; Miyoshi et al., 2016; Parker et al., 2005), we used both substrates containing a 5’ phosphorylate (5P) group and a 5’ hydroxyl group (5OH), respectively. SiAgo showed weak but apparent binding ability to RNA and the preferred substrate is 5P-RNA (Figure S5A). Considering SiAgo and SiAga1 form a complex, we further analyzed the nucleic acid binding ability of the SiAgo-Aga1 complex. The complex showed higher affinity to all tested nucleic acids, compared to SiAgo alone (Figure 6A and B), indicating that SiAga1 assists SiAgo to bind nucleic acid. However, SiAga1 itself did not show detectable binding affinity to any tested nucleic acids (Figure S5B). In addition, the SiAgo-Aga1 complex still preferred 5P-RNA substrates, suggesting that the SiAgo-Aga1 complex uses 5P-RNA as the guide strand (Figure 6A and B).

**Figure 6.**
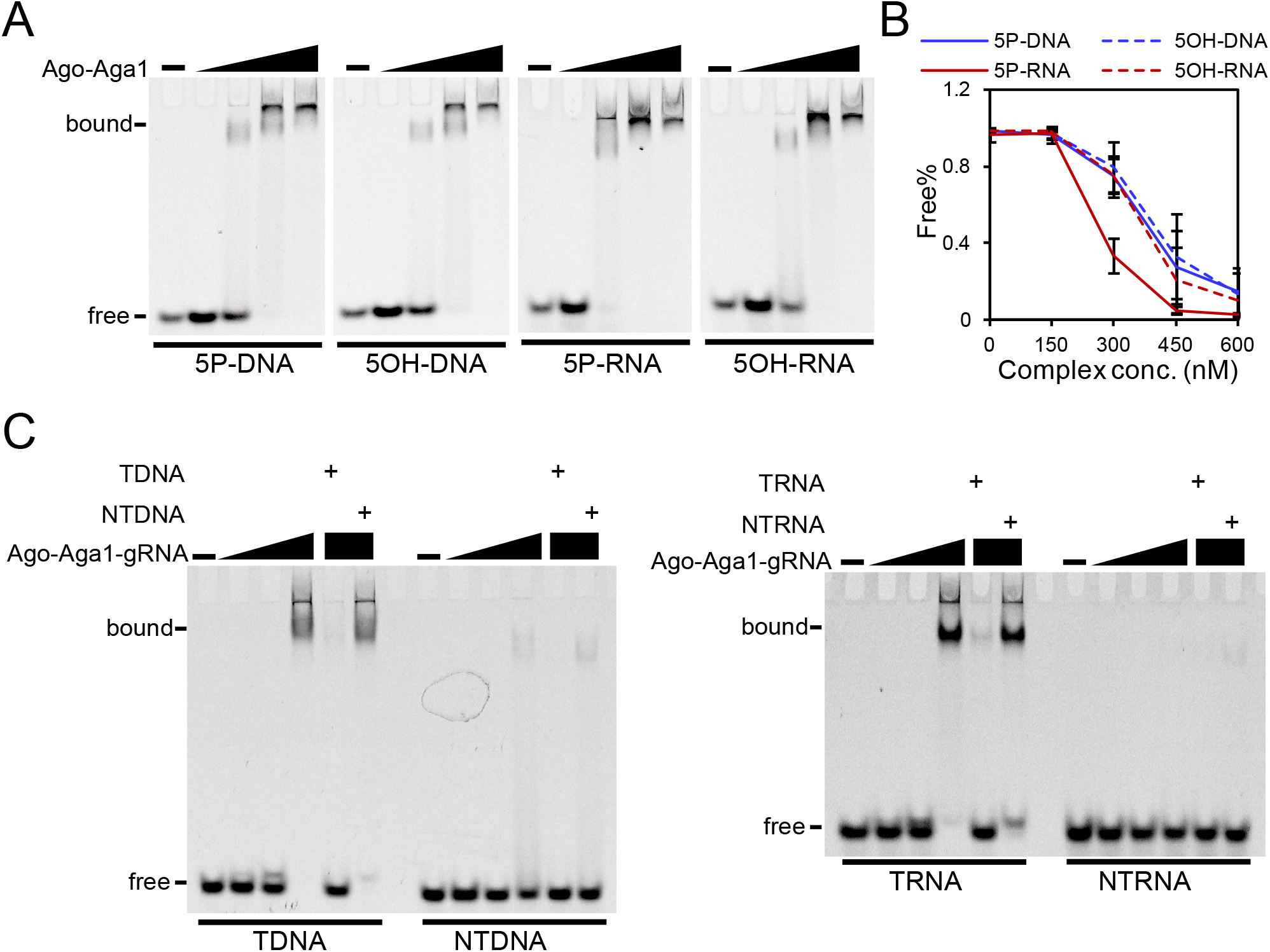
Nucleic acid binding and recognition by SiAgo and SiAga1 complex. (A) Nucleic acid binding of SiAgo-Aga1 complex. The different substrates are indicated below the panels. The concentrations of SiAgo-Aga1 complex were 0, 150, 300, 450, 600 nM, respectively. (B) Quantification of the free substrates from (A). The data show means of three independent replicates. Error bars indicate the standard deviations. (C) Nucleic acid recognition of SiAgo-Aga1 complex preloading with 5P-RNA. SiAgo-Aga1 complex (400 nM) was incubated with 100 nM 5P-RNA. Then, the mixture was serially diluted and further incubated with 50 nM FAM-labeled target DNA (TDNA) and non-target DNA (NTDNA) (left panel), or target RNA (TRNA) and non-target RNA (NTRNA) (right panel). Unlabeled target and non-target nucleic acids were also supplemented up to 1 μM as competitors, as indicated.

Next, we analyzed whether the SiAgo-Aga1 complex possesses guide RNA-directed target nucleic acid recognition ability. The complex was incubated with 5P-RNA at the ratio of 4:1, when most RNA was associated with the complex. Then, the ternary complex was further incubated with nucleic acids that are complementary to guide RNA (TDNA and TRNA) or not (NTDNA and NTRNA). The results show that only the complementary DNA and RNA were efficiently bound by the ternary complex (Figure 6C). Further, addition of 20-fold unlabeled complementary competitors abolished the binding to target nucleic acids, while non-complementary competitors exhibited little effects on the target binding (Figure 6C). The data show that preloading of 5P-RNA endows SiAgo-Aga1 complex with the specific target recognition ability.

### MID domain is important for both Abi induction and target recognition

The MID domain of pAgos is implicated in binding to the 5’ end of the guide. Mutation of the conserved residues of MID domain impairs the guide binding ability and the target silence or cleavage activity (Ma et al., 2005; Miyoshi et al., 2016; Willkomm et al., 2017). SiAgo harbors the conserved MID domain residues as other pAgos (Figure S1C). To analyze whether mutation of the conserved residues would affect target recognition of SiAgo-Aga1 complex, we purified SiAgo-Aga1 complexes carrying a point mutation within the MID domain, either K142A (M1) or K183A (M2), respectively. EMSA analysis indicated that the wild type and mutated complexes showed similar affinity to the 5P-RNA guide (Figure S5C). Next, the complexes were preloaded with guide RNA and analyzed for their affinity to target DNA. The mutated complex still bound target DNA, but the binding resulted in two bands, one of which was similar to that of the wild type complex, while the other band migrated faster in the gel, indicative of a different form of the target-binding complex (Figure 7A). In addition, a band representing the duplex of target DNA and guide RNA was observed for the mutated complexes in the presence of 20-fold non-complementary competitor (NTDNA), suggesting that guide RNA has been released from the mutated complexes. To further explore this phenomenon, we preloaded the complexes with FAM-labeled guide RNA and incubated the ternary complexes with complementary DNA (TDNA), non-complementary DNA (NTDNA), or both. Incubation with the DNA resulted in release of guide RNA from the complexes, and more RNA was released from the mutated complexes than the wild type complex (Figure 7B and C), indicating that the conserved residues play a role in stabilizing the binding to guide RNA.

**Figure 7.**
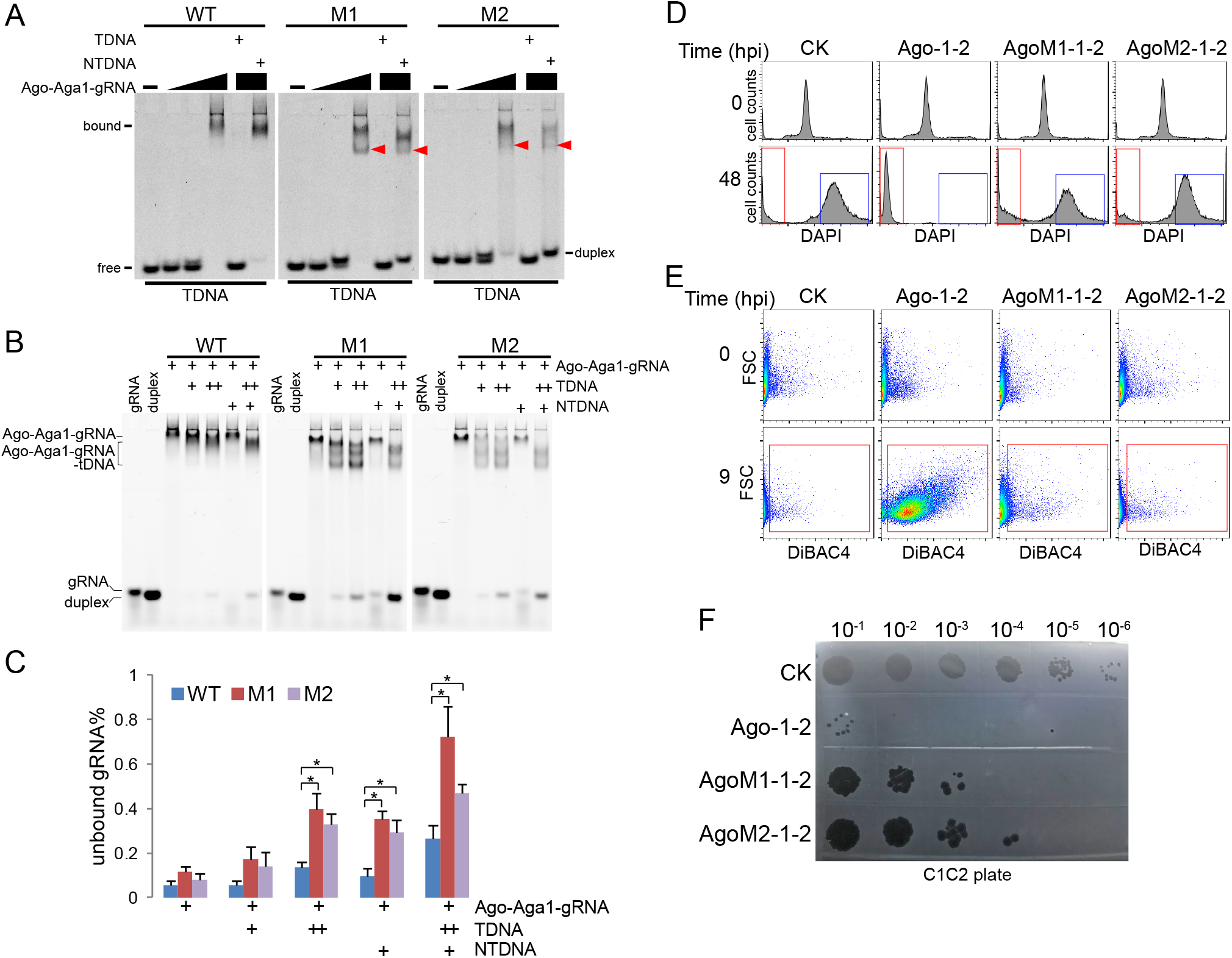
Functional characterization of the conserved MID domain motif. (A) Target DNA binding of the wild type (WT) and mutated Ago-Aga1-gRNA ternary complexes. The ternary complexes were generated by incubating WT and mutated SiAgo-Aga1 complexes (400 nM) with 100 nM 5P-RNA. Then, target DNA was incubated with a gradient of the ternary complexes. WT: wild type; M1: Ago-K142A; M2: Ago-K183A. Non-labeled competitors were supplemented at 1 μM as indicated. “duplex” represents the duplex of 5P-RNA and target DNA. Red arrow indicates a band only observed in gel of the mutated complexes. (B) Release of guide RNA from WT and mutated SiAgo-Aga1 complexes. FAM-labeled guide RNA was bound by WT and mutated SiAgo-Aga1 complexes at first. Then, the ternary complex was incubated with water, target DNA (+: 100 nM; ++: 200 nM), non-target DNA (1 μM) or both. Free guide RNA and RNA/DNA duplex were also loaded onto the gels as markers. (C) Quantification of the unbound RNA from (B). Data shown are means of three independent repeats ± standard deviation. *: P<0.05. (D) Membrane depolarization of the cultures expressing WT or mutated SiAgo systems at 0 and 9 h post SMV1 infection. (E) DNA content distributions of the cultures expressing WT or mutated SiAgo systems at 0 and 48 h post SMV1 infection. (F) Viral titers of the cultures expressing WT or mutated SiAgo systems at 48 hpi.

Next, we analyzed whether the conserved MID residues are important for Abi induction of the SiAgo system. We therefore constructed strains expressing the SiAgo system carrying the K142A and K183A mutations of SiAgo and analyzed the response of the strains to SMV1 infection. The mutated SiAgo system failed to induce DiBAC_4_(3)-positive cells and resulted in similar DNA content distributions to the strain lacking the system (Figure 7D and E), indicating that mutation of MID domain abolishes the Abi induction ability of the SiAgo system. Subsequent PFU analysis showed that the viral titer of the strains carrying the mutated SiAgo system was about 100 times higher than that of the WT system, but still substantially lower than the strain lacking the system (Figure 7F). The data suggest that the mutated SiAgo system can still moderately impair SMV1 release and/or replication by an unknown mechanism.

## Discussion

In this study, we demonstrate that a short pAgo and its two genetically associated proteins (Aga1 and Aga2), mediate an Abi response that confers robust anti-viral immunity in *S. islandicus*. Abi is a defense strategy employed by many prokaryotic defense systems (Isaev et al., 2021; Lopatina et al., 2020) and is composed of three steps: (1) a sensor module recognizes a cue from an invading virus; (2) the sensor module activates a toxic effector (killer module); (3) the effector module induces cellular dormancy or cell death, hence preempting further viral spread in the population. In our study, the cells containing the SiAgo system sequentially experienced membrane depolarization, loss of genomic DNA and loss of membrane integrity post viral infection. The data suggest that SiAgo system directly induces membrane depolarization, which eventually results in cell death. Notably, *S. islandicus* M164, which naturally carries the SiAgo system, turns into a dormant status post viral infection (Bautista et al., 2015). The status is either reversible with active CRISPR-Cas immunity or results in cell death in the absence of CRISPR-Cas immunity. We cannot exclude the possibility that the SiAgo system may mediate the cellular dormancy in *S. islandicus* M164 and provide immunoprotection in cooperation with CRISPR-Cas systems.

In the SiAgo system, the SiAgo-associated protein SiAga2, which is a membrane-associated protein, acts as the killer effector. Membrane proteins are widely found in CBASS (Millman et al., 2020b), retrons (Mestre et al., 2020; Millman et al., 2020a), type III CRISPR-Cas (Shah et al., 2019; Shmakov et al., 2018), Thoreis, Zorya, Kiwa, etc. (Doron et al., 2018) and are predicted to function as toxic effectors. Notably, in addition to SiAga2, the gene contexts of pAgos encode other membrane proteins as well (Ryazansky et al., 2018). Previous bioinformatic analyses show that SIR2 and TIR domains, which are recently found to be implicated in Abi (Ofir et al., 2021), are also found in association with pAgos, especially short pAgos (Ryazansky et al., 2018). Conceivably, pAgo systems could rely on these putative ancillary effector proteins (or domains) to mediate immunity, highlighting Abi as a common strategy for a broad diversity of pAgo systems, especially those lacking intrinsic nuclease activity.

Our findings demonstrate that SiAga2 mediates cell death via membrane depolarization, a phenomenon that has been observed for other Abi systems containing toxic membrane effectors. These effectors are believed to form ion channels or pores that allow free flux of ions or other cellular contents across the cell membrane and result in loss of membrane potential (Duncan-Lowey et al., 2021; Durmaz and Klaenhammer, 2007; Parma et al., 1992). Intriguingly, a similar strategy is also employed by eukaryotes to mediate programmed cell death, including apoptosis, necroptosis and pyroptosis (Ly et al., 2003; Shi et al., 2017; Wang et al., 2014). Moreover, SiAga2 relies on its binding to the anionic head groups of phospholipids to mediate cell death. The characteristic is also shared by many mammal cell-suicide effectors, such as gasdermin, a pore-forming protein which mediates pyroptosis in response to microbial infection (Ding et al., 2016; Liu et al., 2016), and MLKL, a necroptosis effector (Dondelinger et al., 2014; Wang et al., 2014). In addition, SiAga2 forms large oligomers, which is a common feature for pore-forming proteins (Mesa-Galloso et al., 2021). These similarities reveal an unprecedented parallel between archaeal and mammalian cell-suicide immune pathways. Notably, gasdermins have been recently identified in bacteria and shown to form toxic membrane pores (Johnson et al., 2021), further highlighting membrane disruption-mediated cell death as a common immune response mechanism across the three domains of life.

The present study raises a question: what is the native target of SiAga2? *In vitro*, SiAga2 exhibits the highest affinity to PA and PI(n)P, the archaeal counterparts of which are 2,3-di-O-geranylgeranylglyceryl phosphate (DGGGP) and archaetidylinositol phosphate (AIP), respectively. DGGGP and AIP are not the inherent components of the cell membrane but intermediates in lipid synthesis (Jensen et al., 2015; Rastadter et al., 2020). A possible scenario is that SiAga2 binds the intermediates and retains them from further lipid synthesis reactions. In addition, cardiolipin and PS, the less affinitive substrate of SiAga2, only have their counterparts in euryarchaea instead of crenarchaea. Together, the results imply that DGGGP and AIP could serve the potential targets of SiAga2 *in vivo*. Further investigations of the interaction between SiAga2 and its targets are required to provide insights into the membrane disruption mechanisms of this unique archaeal toxic effector.

We demonstrate that SiAgo and SiAga1 form a stable complex that exhibits RNA-guided nucleic acid recognition ability. SiAga1 significantly contributes to nucleic acid binding, suggesting that it may complement the loss of the N-terminal and PAZ domains in SiAgo, which function in guide and target binding in long pAgos (Kaya et al., 2016; Ma et al., 2004; Song et al., 2004). Other short pAgos have been found to be fused or associated with APAZ domain and/or DNA binding domains (Ryazansky et al., 2018), which might also contribute to guide and target binding.

The MID domain of pAgos functions in interacting with the 5’-end of guides (Ma et al., 2005; Miyoshi et al., 2016; Willkomm et al., 2017). In the SiAgo-Aga1 complex, the conserved residues in the MID domain can stabilize the association of guide RNA with the complex, and thus facilitate target binding. Moreover, we show that the conserved residues are essential for mediating Abi, indicative of the importance of nucleic acid recognition in the process. In addition, the SiAgo-Aga1 complex interacts with SiAga2 and can enhance its toxicity. Based on these results, we propose a model that explains how the SiAgo system mediates Abi (Figure 8): the SiAgo-Aga1 complex binds to 5P-RNA derived from viral mRNA as guides to recognize complementary viral nucleic acid targets; then the quaternary complex activates SiAga2 to induce membrane depolarization and execute cell death. The SiAgo-Aga1 complex may discriminate viral nucleic acids in a similar way to RsAgo (Olovnikov et al., 2013), a long-B pAgo that also uses 5P-RNA guides. As proposed for RsAgo, during the virus life cycle, viral genes are highly expressed and the genome is replicated at high rates, thus generating abundant mRNA and single-strand DNA that provide guides and targets for the SiAgo.

**Figure 8.**
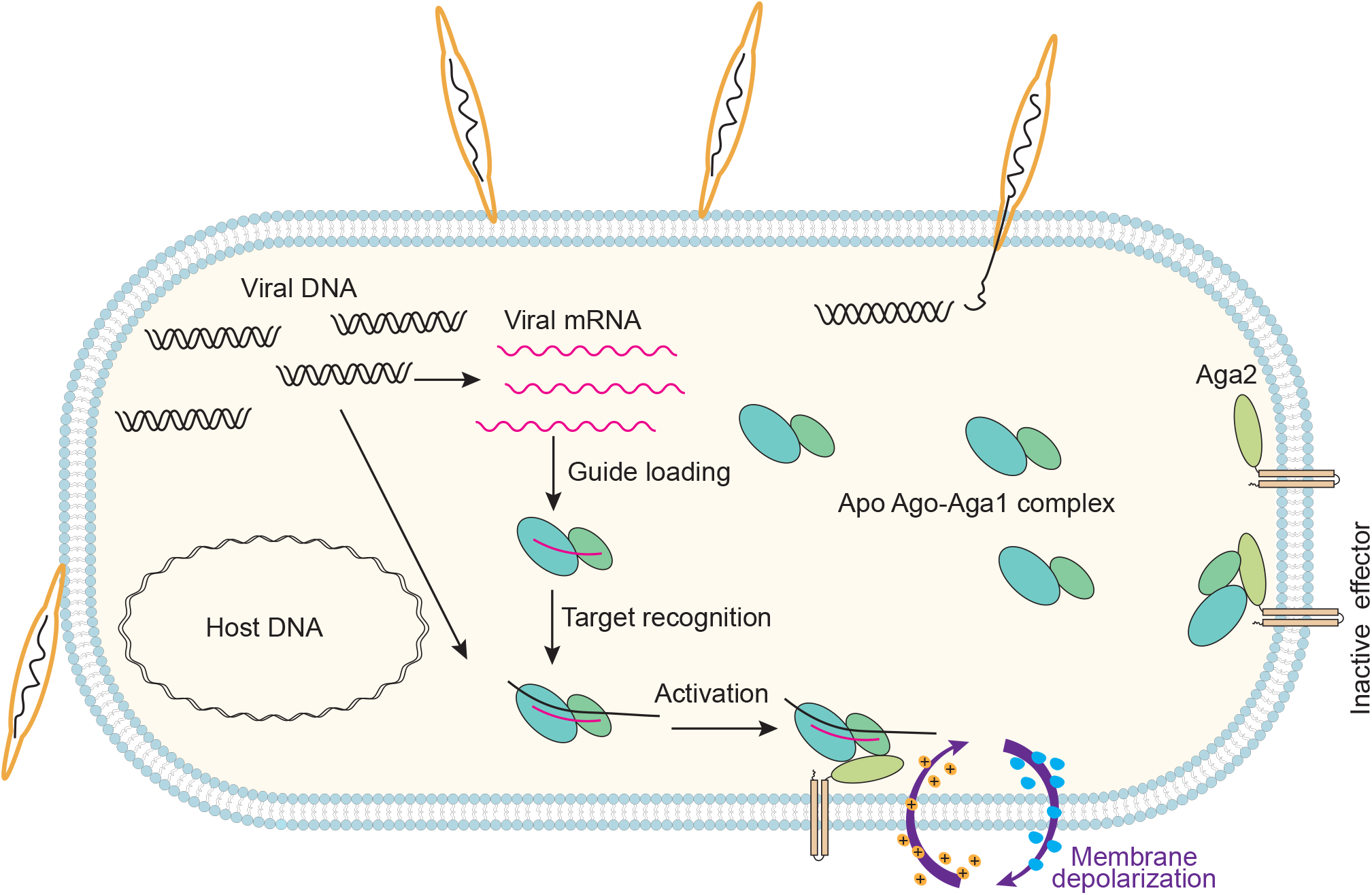
Model for the antiviral immune response of SiAgo system. Prior to viral infection, apo SiAgo-Aga1 complex resides in the cytoplasm or is recruited to SiAga2 on the membrane without triggering it. During viral infection, high transcription of viral genes and fast replication of the viral genome provide abundant RNA and DNA substrates for the SiAgo-Aga1 complex to obtain guides, and also for the guide-loaded complex to search targets. Target binding leads to the activation of SiAga2, probably through direct interaction, which then induces the death of infected cells through membrane depolarization.

Ago and CRISPR-Cas are, to date, the only two sequence-dependent nucleic acid sensing defense systems. While the consequence of nucleic acid sensing is usually nucleic acid degradation (Abudayyeh et al., 2016; Gasiunas et al., 2012; Jinek et al., 2012; Kazlauskiene et al., 2017; Niewoehner et al., 2017; Swarts et al., 2014a; Westra et al., 2012), diverse membrane proteins and other potential toxic effectors are usually found in association with these systems (Ryazansky et al., 2018; Shah et al., 2019; Shmakov et al., 2018; van Beljouw et al., 2021). Our work, together with a recent study on type VI-B2 CRISPR-Cas system (VanderWal et al., 2021), highlights that such association represents a general yet unprecedented defense strategy that leverages nucleic-acid sensing to trigger downstream toxic membrane effectors to confer immunity via cell death.

## Acknowledgments

The research was supported by National Key Research and Development program of China (2019YFA0906400), National Science Foundation of China (Grant No. 31970545), Fundamental Research Funds for Central Universities (Grant No. 2662020SKPY001) and Huazhong Agricultural University Scientific & Technological Self-innovation Foundation. R.P-R. was financed by the Lundbeck Foundation (Lundbeckfonden), postdoc grant R347-2020-2346. S.A.S. is a recipient of a Novo Nordisk Foundation project grant in basic bioscience [NNF18OC0052965]. C.Z. and R.W. were supported by U.S National Science Foundation under IOS grant award no. 1656869. We thank the core facilities of Center for Protein Research (CPR) and Experimental Teaching Center of Bioengineering at Huazhong Agricultural University for technical support.

## Author contributions

Z.Z., Y.C. and Z.H. conducted the experiments. S.A.S. and R.P-R. performed bioinformatics analysis. F.Z. and C.W. predicted and analyzed the structures. C.Z., R.W. and Q.S. provided archaeal materials. R.P-R., C.Z., R.W. and Q.S. critically commented the draft. W.H. acquired the funding, supervised the work and wrote the original draft. All authors contributed to review and editing.

## Competing interests

The authors declare no competing interests.

## STAR METHODS

## RESOURCE AVAILABILITY

### Lead Contact

Further information and requests for resources and reagents should be directed to and will be fulfilled by the Lead Contact, Wenyuan Han.

### Materials Availability

Plasmids, strains and other unique reagents generated in this study are available upon request.

### Data and Code Availability

This study did not generate any unique datasets or code.

## EXPERIMENTAL MODELS AND SUBJECT DETAILS

*S. islandicus* E233S1 (Deng et al., 2009) was used as the genetic host to express the SiAgo system. The strain was grown at 78 °C in SCVU medium as indicated previously (basic salts medium supplemented with 0.2% (w/v) sucrose, 0.2% (w/v) casamino acids, 0.2% (w/v) uracil and a vitamin mixture) (Zhao et al., 2021). The strains carrying a pSeSD-derived expression plasmid were grown in SCV medium lacking uracil. STV medium in which casamino acids were replaced with tryptone was used for large scale cultivation. Arabinose was supplemented at 0.2% (w/v) to induce expression of the proteins of interest. SMV1 (Sulfolobus Monocaudavirus 1) (Uldahl et al., 2016) was used to analyze the immune functions of the SiAgo system and its variants. *S. islandicus* C1C2 (Gudbergsdottir et al., 2011) was used for virus propagation and plaque forming unit (PFU) assay. *Escherichia coli* DH5a and BL21 (DE3) were routinely grown in LB medium and used for plasmid cloning and protein expression, respectively.

## METHOD DETAILS

### Construction of *E. coli* expression plasmids and strains

To express the proteins in *E. coli*, the coding sequences of SiAgo and SiAga1 were amplified from the genomic DNA of *S. islandicus* M164 by PCR using the primers listed in Table S4. The gene fragments were inserted in pET30aN between the NheI and NotI restriction sites, such that the encoded proteins have an N-terminal 6×His tag. The pET30aN vector was constructed by inserting an NheI site into pET30a to increase the compatibility of the restriction sites between pSeSD and pET30a. Since the coding sequence of SiAga2 that was amplified from the genomic DNA resulted in poor protein expression in *E. coli*, we ordered a synthetic, codon-optimized sequence of SiAga2 from Tingke (Beijing, China) (Table S3). To facilitate protein expression and stability, the synthesized gene was fused with the coding sequence of the maltose-binding protein (MBP). First, the optimized coding sequence of SiAga2 was inserted into pMAL-c5x between the NdeI and NotI sites; then, the fused gene fragment of MBP-Aga2 was amplified by PCR and inserted into pCDFDuet-1 between BamHI and NotI, yielding pCDFDuet-1-Aga2 encoding the His-MBPAga2 fusion protein. To express SiAga2ΔC, a truncated version that lacks the transmembrane region, the coding sequence was amplified from the pCDFDuet-1-Aga2 plasmid and inserted into pET30aN between the NheI and XhoI sites. The resulting plasmid constructs, i.e. pET30aN-Ago, pET30aN-Aga1, pCDFDuet-1-Aga2 and pET30aN-Aga2ΔC, were transformed into *E. coli* BL21(DE3) to generate strains for protein expression.

To perform the co-expression assay, we firstly constructed the plasmids expressing the His-tag free (HF) SiAga1 and SiAga2. The *aga1* gene was inserted into pET21d between Nde1 and Xho1, while the fused gene of *mbpaga2* was inserted in pCDFDuet-1 between NdeI and XhoI, generating pET21d-Aga1(HF) and pCDFDuet-1-Aga2(HF), respectively. pCDFDuet-1-MBP(HF), which encodes His-tag free MBP, was also constructed in a similar way. To obtain the co-expression strains, either two or three plasmids were simultaneously transformed into *E. coli* BL21(DE3). The obtained co-expression strains include those carrying pET30aN-Ago + pET21d-Aga1(HF), pET30aN-Ago + pCDFDuet-1-Aga2(HF), pET30aN-Aga1 + pCDFDuet-1-Aga2(HF), pET30aN-Ago + pET21d-Aga1(HF) + pCDFDuet-1-Aga2(HF), pET30aN-Ago + pET21d-Aga1(HF) + pCDFDuet-1-MBP(HF).

To express the mutated SiAgo and SiAga2ΔC, overlapping PCR was performed to generate site mutations within the coding sequences, and the mutated genes were inserted in pET30aN as indicated above. The plasmids encoding mutated SiAgo were also co-transformed with pET21d-Aga1(HF) to express mutated SiAgo-Aga1 complexes. All the primers were synthesized by Tingke (Beijing, China). All the plasmids were confirmed by sequencing before transformation. The plasmids are listed in Table S1.

### Construction of *Sulfolobus* expression plasmids and strains

The pSeSD shuttle vector (Peng et al., 2012) was used to construct the plasmids expressing Ago and its associated proteins in *S. islandicus* E233S1 (Table S1). To construct them, the coding sequences of SiAgo, SiAga1, SiAga2, and SiAga1+Aga2 were amplified from the genomic DNA of *S. islandicus* M164. The coding sequences of SiAga1, SiAga2, and SiAga1+Aga2 were inserted between NdeI and NotI in pSeSD, yielding pSeSD-Aga1, pSeSD-Aga2 and pSeSD-Aga1-Aga2, while the *ago* gene was inserted between NheI and NotI, resulting in pSeSD-Ago. To co-express SiAgo and its associated proteins, the expression cassette of SiAgo from the pSeSD-Ago plasmid was amplified using the primers Aras-F_SmaI and Ago-R_NotI with the plasmid as template and inserted in pSeSD-Aga1, pSeSD-Aga2 and pSeSD-Aga1-Aga2 between NotI and SmaI respectively, generating pSeSD-Ago-Aga1, pSeSD-Ago-Aga2 and pSeSD-Ago-1-2. When co-expressed, only SiAgo carries an N-terminal His tag. The plasmids expressing mutated SiAgo and SiAga2 were constructed in a similar way, except that the wild type coding sequences were replaced by those carrying the indicated mutations. Then, the plasmids were transformed into *S. islandicus* E233S1 to generate the corresponding expression strains (Table S2). The detailed procedures of *S. islandicus* E233S1 cultivation and transformation were described previously (Zhao et al., 2021). All the primers were synthesized by Tingke (Beijing, China). All the plasmids were confirmed by sequencing before transformation.

### Protein expression and purification

To express the proteins from *E. coli* BL21(DE3), the strains carrying the indicated plasmids were grown in LB medium containing the corresponding antibiotics. At an optical density of ∼1.0, protein expression was induced with 0.5 mM IPTG at 18 °C for 18 h. To express the proteins from *S. islandicus* E233S1, the strain carrying corresponding plasmids was grown to an optical density of ∼0.6 in STV medium and then protein expression was induced by 0.2% (w/v) arabinose for 24 h. Then, the cells were collected by centrifugation at 7000 g for 10 min.

The cell mass was resuspended in 50 ml of lysis buffer (20 mM HEPES pH 7.5, 20 mM imidazole, 500 mM NaCl) and stored at -80 °C until protein purification.

The purification procedure for all the protein and protein complexes was similar. The common steps include cell extract preparation, Ni-NTA affinity chromatography (NAC), anion exchange chromatography (AEC) and size exclusion chromatography (SEC). When full length SiAga2 was purified, 1% n-Dodecyl-β-D-Maltopyranoside (DDM) (RHAWN, Shanghai, China) was used to treat the cell extract at 4 °C for 12 h and the buffers used for NAC, AEC and SEC contained 0.02% DDM.

To prepare cell extracts, the cells were disrupted by French press and the lysate was subjected to centrifugation at 13000 g for 40 min to remove cell debris. Before centrifugation, the cell lysate was treated with 1% DDM if full length SiAga2 was purified. Then, the cell extract was loaded onto Ni-NTA agarose resin columns (Cytiva, Marlborough, MA, USA). After the column was washed with lysis buffer containing 60 mM imidazole, His-tagged proteins were eluted using a lysis buffer containing 300 mM imidazole. Then, the elution fractions were concentrated employing an ultra-centrifugal filter (Eppendorf, Hamburg, Germany) and then diluted for 30-fold with Buffer A (25 mM Tris-HCl, pH 8.0). The diluted samples were loaded onto a 5 mL Q FF column (Cytiva, Marlborough, MA, USA) and the proteins were eluted using a 35 mL linear gradient of 0-1 M NaCl. The fractions containing target proteins were concentrated again and loaded onto a Superdex 200 column (Cytiva, Marlborough, MA, USA). Finally, the proteins were eluted with BufferC (20 mM Tris-HCl pH 7.5, 250 mM NaCl) and analyzed by SDS-PAGE.

### Flow cytometry

The cellular DNA content, membrane polarity and membrane permeability were analyzed with flow cytometry. The cell samples were prepared following procedures previously established in our group with some modifications (Han et al., 2017). Specifically, the cells for cellular DNA content analysis were fixed with 70% ethanol at 4 °C for at least 12 h, and then washed with 1 mL of washing buffer (10 mM Tris-HCl, pH 7.5, 10 mM MgCl_2_). The cells were collected again and resuspended in 30 μL washing buffer. DAPI (Thermo Scientific, Waltham, MA, USA) was added into the cell suspension to a final concentration of 3.3 μg/mL and the cells were stained for 30 min on ice in darkness. Then, the cell suspensions were diluted in 1 mL and loaded onto a cytoflex-LX flow cytometer (Beckman Coulter, Brea, CA, USA) with a 375 nm laser. A dataset of at least 40,000 cells was recorded for each sample, including fluorescence signal at 450 nm, FSC (forward scattered light), and SSC (side scattered light). At last, the data were analyzed and visualized by FlowJo v.10 (BD Biosciences, Franklin Lakes, NJ, USA).

The cells for the analyses of membrane polarity and membrane permeability were collected from 0.1 mL culture, washed with fresh medium, and resuspended in 50 μL fresh medium. Then, 0.5 μL of dye mix containing SYTO 9 and propidium iodide (PI) in the ratio 1:1 (LIVE/DEAD BacLight bacterial viability kit) (Thermo Scientific, Waltham, MA, USA) was supplemented to stain the cells for membrane permeability analysis, while DiBAC_4_(3) (Sigma-Aldrich, St. Louis, MO, USA) was added up to the concentration of 0.5 µg/ml for membrane polarity analysis. The cells were stained in darkness for 15 min at room temperature and analyzed by Cytoflex-LX flow cytometer with a 488 nm laser. For the SYTO9/PI-stained cells, fluorescence signal at 525 nm (green, SYTO9) and 610 nm (red, PI) was analysed, while the green signal was measured after DiBAC straining. The data were also analyzed and visualized by FlowJo v.10.

### Virus propagation

A sensitive strain, *S. islandicus* C1C2 (Gudbergsdottir et al., 2011), was grown at exponential phase for at least 72 h. At an optical density at 600 nm (OD_600_) of ∼0.2, 100 mL of the culture was infected with SMV1 at a MOI of <0.1. At 48 h post infection (hpi), the cells were removed by centrifugation at 8000 g for 10 min and the virus particles in the supernatant were concentrated by ultrafiltration using 1,000,000-molecular-weight-cutoff (MWCO) centrifugal filter units (Sigma-Aldrich, St. Louis, MO, USA). The concentrated virus particles were then dissolved in viral storage buffer containing 20 mM Tris-HCl, pH=7.0, 20% glycerol and stored at 4 °C before use.

### Plaque forming unit assay

SCVU plates were prepared before the assay (Zhao et al., 2021). To determine viral titer in the storage buffer, the storage buffer was serially diluted and mixed with 4 mL fresh *S. islandicus* C1C2 culture (OD_600_ around 0.2). The mixture was preheated at 75 °C and further mixed with an equal volume of preheated 0.4% Gelzan CM (Duchefa-biochemie, Haarlem, Netherlands). Then, the mixture was spread onto pre-warmed SCVU plates. Plaques were counted after the plates had been incubated at 75 °C for two days.

To perform the drop plaque assay, C1C2 plates were firstly prepared by spreading the mixture of 4 mL preheated *S. islandicus* C1C2 culture (OD_600_ around 0.2) and 4 mL preheated 0.4% Gelzan CM onto SCVU plates. Then, cells were removed from the SMV1-infected cultures and the supernatant was serially diluted. Ten μL of the diluted supernatant (from 10^−1^ to 10^−6^) was dropped on the C1C2 plates. Pictures of the plates were taken after incubation at 75 °C for two days.

### Viral infection and sampling

The strains were grown at exponential phase for at least 72 h. At an optical density of ∼0.1, 50 mL of the cultures were infected with SMV1 at a MOI of ∼2. The MOI was calculated based on the estimation that 1 mL of OD_600_=0.1 culture contains 1×10^8^ cells. At indicated time points, samples were removed from the cultures for analysis. When applicable, the cultured cells were collected and diluted to an optical density of ∼0.1 in fresh medium at 72 hpi.

### Membrane and PIP strip screening assay

Membrane and PIP strips (Echelon Biosciences, Salt Lake City, UT, USA) were firstly blocked overnight at 4 °C in a blocking buffer (20mM Tris-HCl pH 8.0, 100 mM NaCl, 0.1% Tween 40, 3% BSA), then incubated with a mixture of SiAga2ΔC (100 ng/mL) and anti-SiAga2 serum (1:10000 dilution) in the blocking buffer for 2 h at room temperature. Then, the membrane was washed 3 times with washing buffer (20mM Tris-HCl pH 8.0, 100 mM NaCl, 0.1% Tween 40) for 10 min. Next, the membrane was incubated with blocking buffer containing HRP-conjugated secondary antibody (1:40000 dilution) (Goat Anti-Rabbit IgG) (Abbkine, Wuhan, China) for 30 min at room temperature, followed by another 3 times’ washing step with washing buffer (20mM Tris-HCl pH 8.0, 100 mM NaCl, 0.1% Tween 40) for 10 min. Finally, the signal was detected by the Clarity Western ECL Substrate (Bio-Rad, Hercules, CA, USA) and the membrane was scanned with Tanon 5200 (Tanon, Shanghai, China). The affinity of SiAga2ΔC mutants was analyzed in the same procedure.

### Electrophoretic mobility shift assay (EMSA)

To analyze the nucleic acid affinity of different proteins, four different 3’-FAM labeled nucleic acid substrates (5P-DNA, 5OH-DNA, 5P-RNA and 5OH-RNA) were incubated with SiAgo, SiAga1 and SiAgo-Aga1 complex, respectively, in a 10-μL mixture at 70 °C for 20 min. The incubation buffer, i.e. EMSA buffer, contained 20 mM Mes pH 6.0 and 5 mM MnCl_2_. The sequences of the substrates are listed in Table S3. The concentration of the substrates was fixed as 100 nM, while the concentrations of proteins varied as indicated in the figure legends. The concentration of SiAgo-Aga1 complex was calculated based on the estimation that the stoichiometry of SiAgo and SiAga1 is 1:1 in the complex. After incubation, the reaction samples were mixed with 4 μL loading dye containing 60% glycerol, 0.1% bromophenol blue and 0.1% xylene cyanol, and loaded onto 8% native polyacrylamide gels. The electrophoresis was performed in 0.5×TB buffer (44.5 mM Tris, 44.5 mM boric acid) at 100 V for 1 h. At last, the fluorescent signal was visualized using a Fujifilm FLA-5100 scanner (Fujifilm Life Science, Japan).

To analyze the target binding ability of the SiAgo-Aga1-gRNA ribonucleoprotein (RNP) complex, 400 nM SiAgo-Aga1 complex was incubated with 100 nM unlabeled 5P-RNA in the EMSA buffer at 70 °C for 20 min. Aliquots of the mixture were diluted 2 and 4 times, respectively. Then, 50 nM of FAM-labeled target and non-target DNA or RNA substrates (TDNA, NTDNA, TRNA and NTRNA) were supplemented into the mixture and further incubated at 70 °C for 20 min. Competitive substrates were also supplemented in aliquots up to a final concentration of 1 μM. At last, the samples were analyzed by native gel electrophoresis and visualized with a Fujifilm scanner. The guide binding and target binding abilities of the mutated SiAgo-Aga1 complexes were analyzed in the same way as the wild type complex.

To analyze the stability of the SiAgo-Aga1-gRNA RNP, 400 nM SiAgo-Aga1 complex was incubated with 100 nM FAM-labeled 5P-RNA in the EMSA buffer at 70 °C for 20 min. Then, aliquots of the mixture were further incubated with water as control, unlabeled target DNA (100 nM and 200 nM), non-target DNA (1 μM), or both, at 70 °C for 20 min. The samples were then analyzed by native gel electrophoresis and visualized with a Fujifilm scanner. The mutated SiAgo-Aga1 complexes were analyzed in the same way as the wild type complex.

### Immunofluorescence microscopy

To analyze the subcellular localization of SiAga2, the strain expressing SiAga2 was transferred into Ara medium. After 24 h, cells from 3 mL cultures were collected by centrifugation at 3000 g for 10 min, washed with PBST buffer (137 mM NaCl, 2.7 mM KCl, 10 mM Na_2_HPO_4_, 2 mM KH_2_PO_4_, 0.05% Tween 40, pH7.6), and resuspended in 300 μL PBST buffer. Cold ethanol was added into the cell solution up to a final concentration of 70% (v/v) to fix the cells for at least 12 h. Then, the cells were collected again, washed with PBST buffer twice, and blocked in PBST buffer containing 2% BSA for 2 h at room temperature. Next, the cells were incubated with anti-SiAga2 serum (1:1000 dilution) in the PBST buffer containing 2% BSA at 4 °C overnight. This was followed by an incubation with secondary antibody Dylight 488 Affinipure Goat anti-Rabbit IgG (H+L)(1:1000 dilution)(Abbkine, Wuhan, China) in the PBST buffer containing 2% BSA at room temperature for 2 h. Before and after each incubation, the cells were washed with PBST buffer for three times. After the last washing, the cells were resuspended in 30 μL PBST containing 3.3 μg/mL DAPI and 10 ng/μL membrane staining dye FM4-64X (Thermo Scientific, Waltham, MA, USA), and incubated for 10 min on ice in darkness. At last, 3 μL of the cell solution was dropped onto a slide and photos were captured with a Nikon A1 HD25 camera (Nikon, Japan).

### Western blot

Proteins from SDS-PAGE gels were transferred onto a PVDF membrane (Bio-Rad, Hercules, CA, USA) using Trans-Blot SD Semi-Dry Transfer Cell (Bio-Rad, Hercules, CA, USA). The membrane was blocked in TBST buffer (50 mM Tris, 100 mM Nacl, 0.05% Tween 40, pH 8.0) containing 6% milk, followed by incubation with anti-SiAga2 serum or anti-SiAgo serum, as indicated. Upon completion of three consecutive washing steps, the membrane was incubated with the secondary antibody (Goat Anti-Rabbit IgG) (Abbkine, Wuhan, China). After removing unspecific binding, the Clarity Western ECL Substrate (Bio-Rad, Hercules, CA, USA) was dropped onto the membrane and the signals were recorded using Tanon 5200 (Tanon, Shanghai, China).

### Bioinformatics analysis

Genetic neighborhood analysis and visualization was performed employing the EFI server (Gerlt, 2017). The dotplot between *S. islandicus* strains M164 and M1425 were drawn using the “nucmer” command from the MUMmer 4 software package (https://github.com/mummer4/mummer). Sequence alignment was carried out using Clustal W, and the results were visualized with Jalview (Waterhouse et al., 2009). HHpred (Soding et al., 2005) was used to identify putative functional domains in M164_1612, M164_1613 and M164_1615. The structure of SiAga2 was predicted by AlpfaFold2 following the published procedures (Jumper et al., 2021). The structural model that shows the highest predicted lDDT (Local Distance Difference Test) was selected for further analysis.

## QUANTIFICATION AND STATISTICAL ANALYSIS

### Growth curves

The optical density at 600 nm was measured for each culture and the values were plotted to time (in hours). The data are either displayed as means of three independent replicates, with error bars representing standard deviations (Figure 1), or shown in each replicate (Figure 2, Figure 5 and Figure S2)

### Flow cytometry

Membrane-depolarized cells, i.e. DiBAC_4_(3)-positive cells, are quantified by FlowJo v.10. The data show means of three independent replicates, with error bars representing standard deviations (Figure 5).

## EMSA

Quantification of labeled substrate in the gels was performed with ImageJ. Error bars show standard deviations of three independent experiments (Figure 6 and Figure 7).

The unpaired t-test was used to calculate the P-value: <0.05 = *; <0.01 = **.

## Notes

### Competing Interest Statement

The authors have declared no competing interest.

