## Supplementary materials for "A short prokaryotic argonaute cooperates with membrane effector to confer antiviral defense"

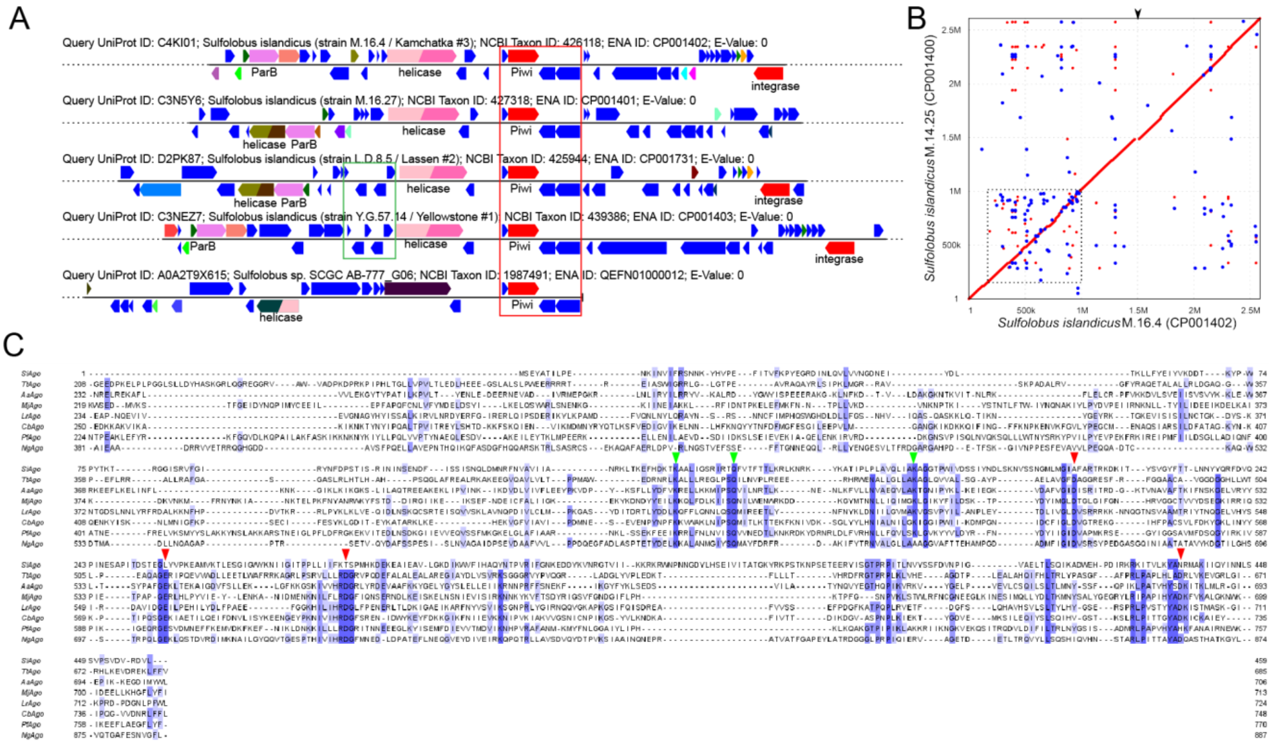


D

| No. | Hit | Name | Probability | E-value | Score |
|---|---|---|---|---|---|
| 1 | 1Z91_A | Organic hydroperoxide resistance transcriptional regulator | 99.2 | 9.9e-10 | 58.41 |
| 2 | 7DVR_A | HTH marR-type domain-containing protein | 99.19 | 4.7e-9 | 56.19 |
| 3 | 6ABT_B | PadR family transcriptional regulator | 99.13 | 1.3e-8 | 50.5 |
| 4 | COG5631 | Predicted transcription regulator, contains HTH domain, MarR family | 99.1 | 1.3e-8 | 56.91 |
| 5 | 2NNN_I | Probable transcriptional regulator | 99.08 | 1.9e-8 | 52.95 |
| 6 | 5HS7_A | HTH-type transcriptional regulator YodB | 99.06 | 4e-8 | 49.81 |

Figure S1. Bioinformatics analysis of *S. islandicus* Ago. Related to Figure 1.

(A) Genome neighborhood analysis of SiAgo using EFI server (https://efi.igb.illinois.edu//efi-gnt/index.php). The red box indicates the invariable four genes that constitute the SiAgo system, while the genes in the green box encode conserved conjugation plasmid proteins.

(B) Dot-plot alignment between *S. islandicus* M164 and *S. islandicus* M1425. Syntenous regions are shown in red, while inversions are blue. The genomic hypervariable region containing most other defence systems is indicated by the dashed rectangle. The region containing the SiAgo system on the M164 genome is indicated by the black arrow in the top.

(C) Sequence alignment of SiAgo with selected long A group pAgos. Only the sequences from the MID and PIWI domains are shown. Green arrows indicate conserved MID domain residues, while red arrows indicate the catalytic residues of PIWI domain, which are mutated in SiAgo.

(D) Searching homologs of M164_1615 using HHpred. Top six hits are listed in the table.

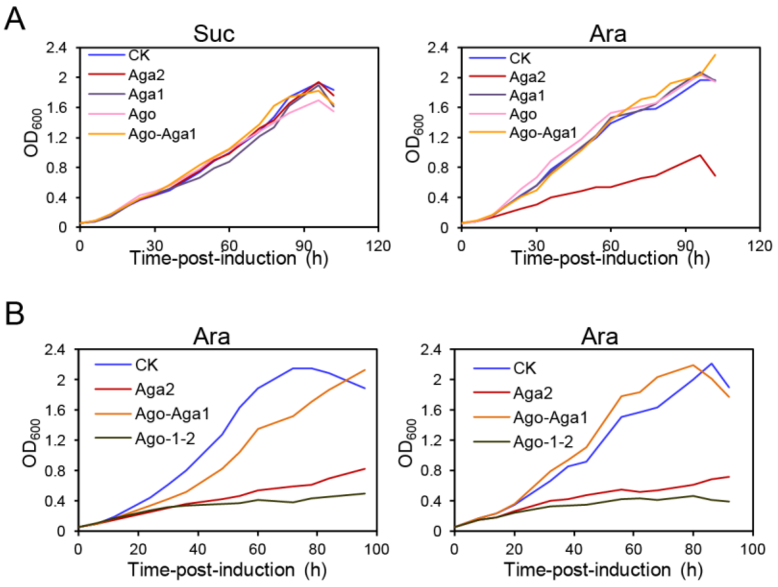

Figure S2. Replicates of growth curves. Related to Figure 2 and 5.

(A) Growth curves of the strains lacking the SiAgo system (CK), or expressing different components of the SiAgo system in sucrose and arabinose medium respectively.

(B) Growth curves of the strains lacking the SiAgo system (CK), or expressing SiAga2, SiAgo+SiAga1 or all the three proteins (Ago-1-2) in arabinose medium. Two independent replicates are shown.

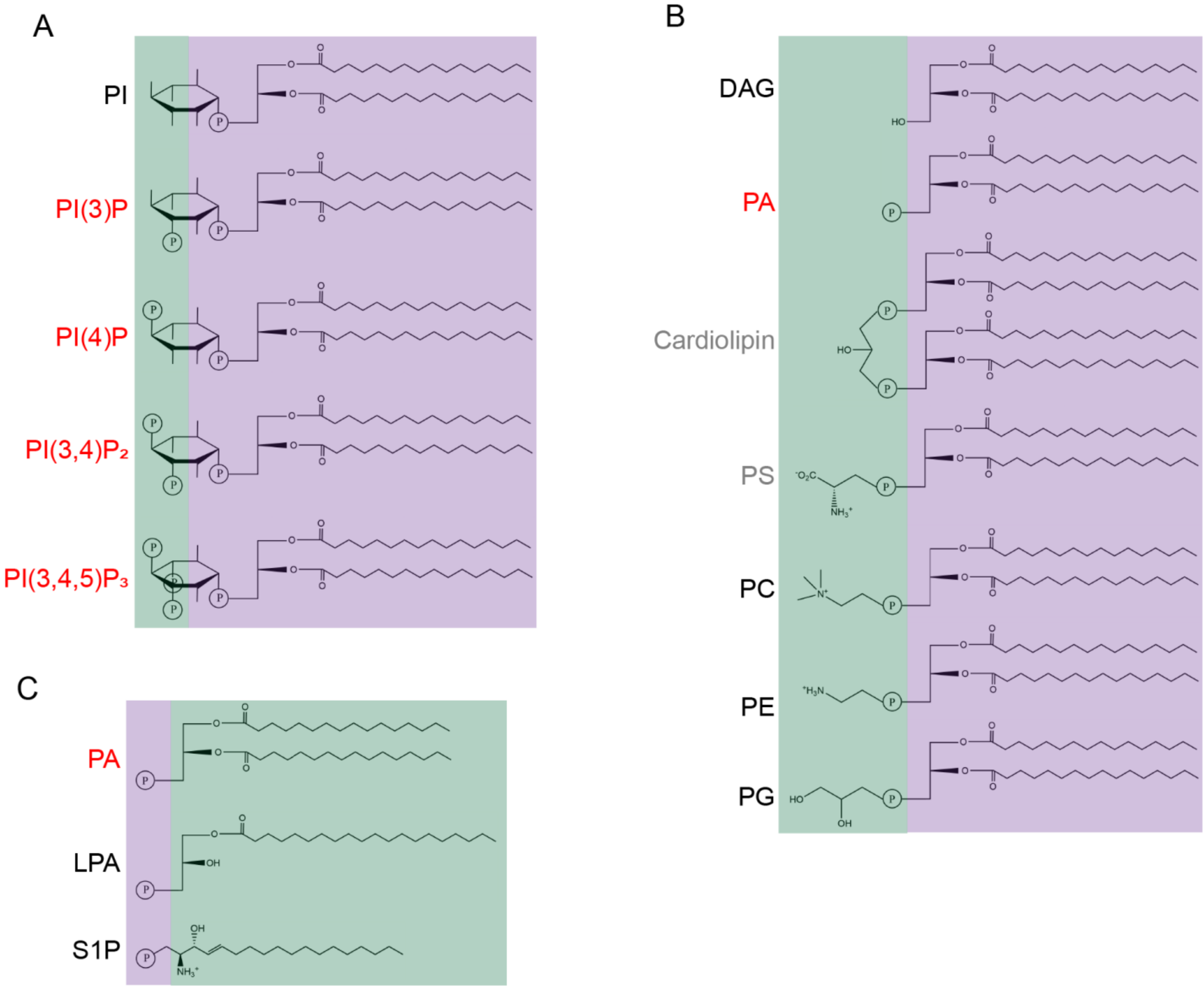


Figure S3. Comparison of affinitive and non-affinitive lipids of SiAga2ΔC. Related to Figure 3.

(A) Comparison of PI and selected PI(n)Ps.

(B) Comparison of DAG, PA, cardiolipin, PS, PE, PC and PG.

(C) Comparison of PA, LPA and S1P.

The common moieties of the lipids are indicated by purple background, while the different moieties are shown in green background. The affinitive lipids are marked in red letters, while the non-affinitive lipids in black letters. PS and cardiolipin, showing lower affinity to SiAga2ΔC, are in grey letters.

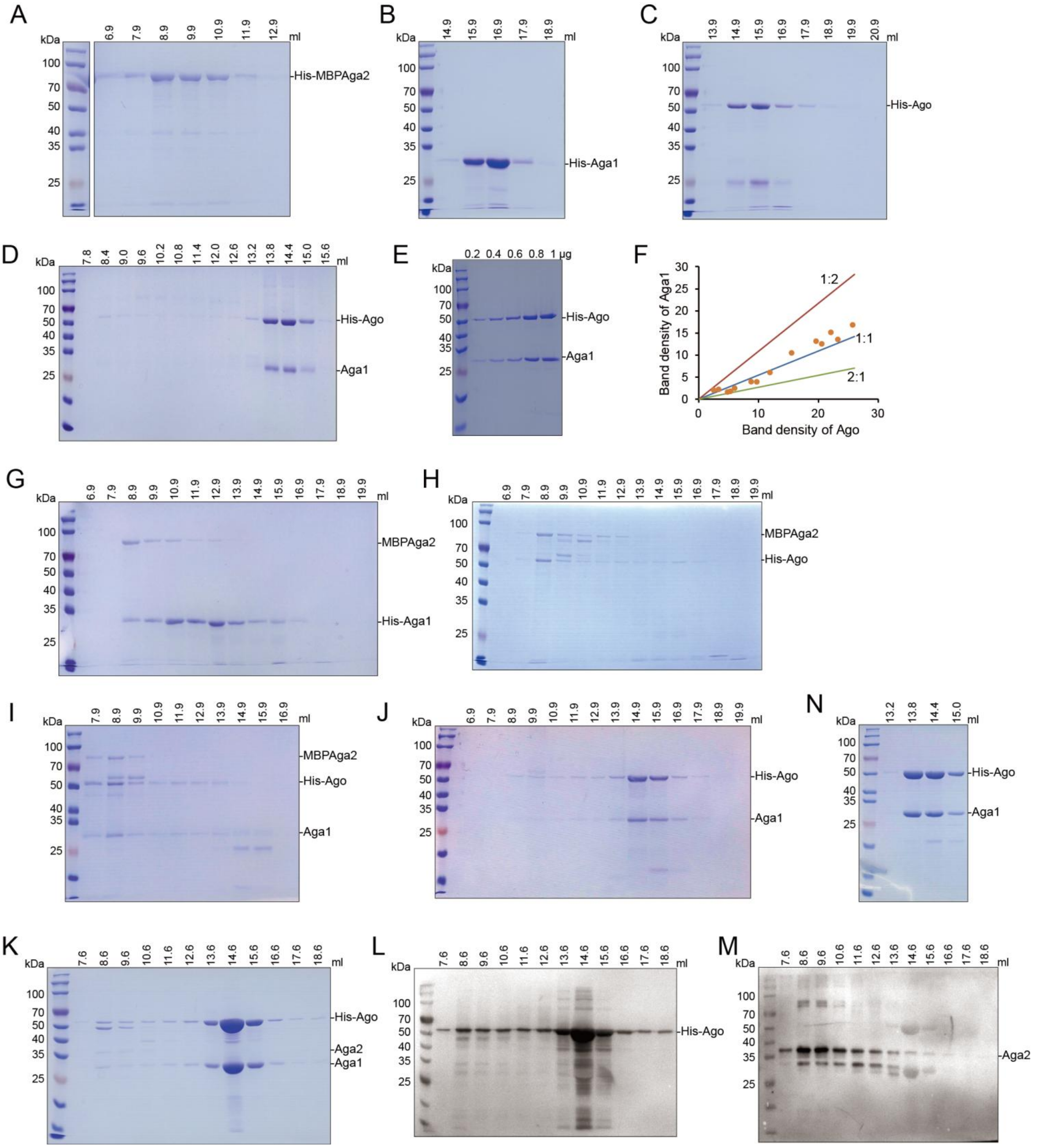


Figure S4. SDS-PAGE and western blot analysis of gel filtration samples after protein purification. Related to Figure 5.

(A)~(C) SDS-PAGE analysis of the gel filtration samples of purified His-MBPAga2, His-Aga1 and His-Ago, respectively.

(D) SDS-PAGE analysis of the gel filtration samples of co-purified His-Ago+Aga1.

(E) A representative gel showing a gradient of SiAgo-Aga1 complex. The amount of the complex loaded was indicated above the gel.

(F) The band density of SiAga1 was plotted to that of SiAgo (orange dots). The three curves show the theoretical relative band density when the stoichometry of SiAgo and SiAga1 is 1:2, 1:1, 2:1, respectively. The dots are well fit with the 1:1 curve.
(G) and (H) SDS-PAGE analysis of the gel filtration samples of co-purified His-Aga1+MBPAga2 (H), and His-Ago+MBPAga2 (F), respectively.
(I) SDS-PAGE analysis of the gel filtration samples of co-purified His-Ago+Aga1+MBPAga2.
(J) SDS-PAGE analysis of the gel filtration samples of co-purified His-Ago+Aga1+MBP.
(K) SDS-PAGE analysis of the gel filtration samples of co-purified His-Ago+Aga1+Aga2 from *Sulfolobus* cells.
(L) and (M) Western blot analysis of samples from (I) using antiserums against SiAgo (J) and SiAga2 (K), respectively.
(N) SDS-PAGE analysis of the gel filtration samples of co-purified His-Ago+Aga1 from *Sulfolobus* cells.

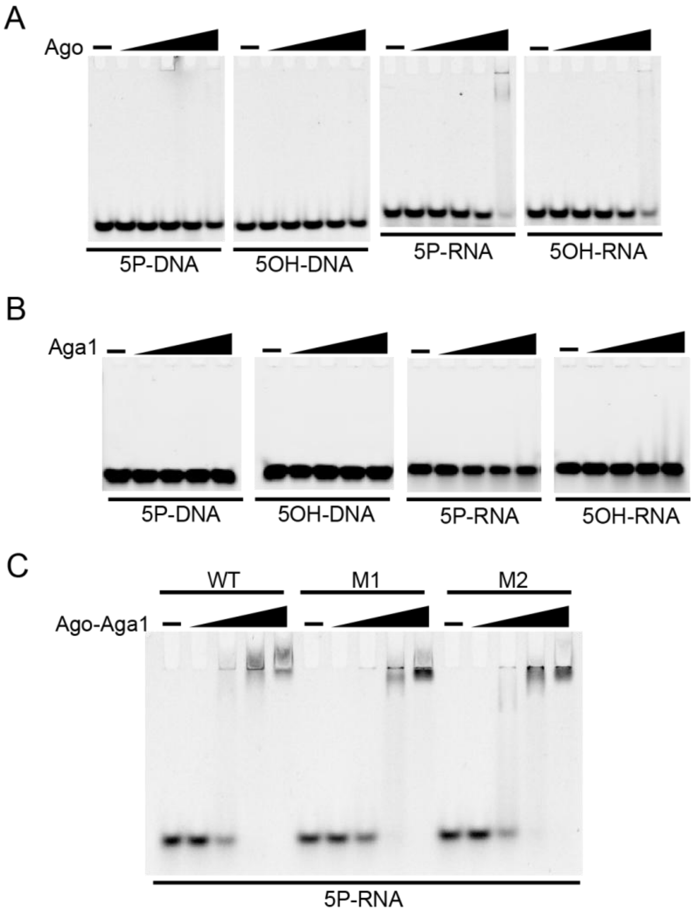


Figure S5. Gel shift analysis of nucleic acids binding. Related to Figure 6 and 7.

(A) The four substrates were incubated with SiAgo gradients (200, 400, 600, 800, 1000 nM) and then analyzed with native PAGE.

(B) The four substrates were incubated with SiAga1 gradients (400, 800, 1200, 1600 nM) and then analyzed with native PAGE.

(C) 5P-RNA was incubated wild type and mutated SiAgo-Aga1 complex gradients (130, 260, 390, 520 nM) and then analyzed with native PAGE.

Table S1. Plasmids constructed in the study

| Plasmids | Description |
|---|---|
| **Plasmids used for protein expression in E.coli** | |
| pET30aN | pET30a carrying an NheI restriction site |
| pET30aN-Ago | To express His-tagged SiAgo in E.coli |
| pET30aN-Aga1 | To express His-tagged SiAga1 in E.coli |
| pMAL-c5x-Aga2 | To generate the coding sequence of MBP-Aga2 |
| pCDFDuet-1-Aga2 | To express His-tagged MBP-fused SiAga2 in E.coli |
| pET30aN-Aga2ΔC | To express His-tagged truncated SiAga2 in E.coli |
| pET21d-Aga1(HF) | To express His-tag free SiAga1 in E.coli |
| pCDFDuet-1-Aga2(HF) | To express His-tag free MBP-fused SiAga2 in E.coli |
| pCDFDuet-1-MBP(HF) | To express His-tag free MBP in E.coli |
| pET30aN-AgoM1 | To express His-tagged SiAgo carrying a K142A mutation in E.coli |
| pET30aN-AgoM2 | To express His-tagged SiAgo carrying a K183A mutation in E.coli |
| pET30aN-Aga2ΔCM1 | To express His-tagged truncated SiAga2 carrying R7A-R8A mutations in E.coli |
| pET30aN-Aga2ΔCM2 | To express His-tagged truncated SiAga2 carrying K12A-K13A mutations in E.coli |
| pET30aN-Aga2ΔCM3 | To express His-tagged truncated SiAga2 carrying R7A-R8A-K12A-K13A mutations in E.coli |
| **Plasmids used for protein expression in S. islandicus** | |
| pSeSD-Ago | Derived from pSeSD; to express His-tagged SiAgo in S. islandicus |
| pSeSD-Aga1 | To express His-tagged SiAga1 in S. islandicus |
| pSeSD-Aga2 | To express His-tagged SiAga2 in S. islandicus |
| pSeSD-Aga1-Aga2 | To express His-tagged SiAga1 and His-tag free SiAga2 in S. islandicus |
| pSeSD-Ago-Aga1 | To express His-tagged SiAgo and His-tag free SiAga1 in S. islandicus |
| pSeSD-Ago-Aga2 | To express His-tagged SiAgo and His-tag free SiAga2 in S. islandicus |
| pSeSD-Ago-1-2 | To express His-tagged SiAgo, His-tag free SiAga1 and His-tag free SiAga2 in S. islandicus |
| pSeSD-Ago-1-2M1 | To express the SiAgo system, with SiAga2 carrying R7A-R8A mutations |
| pSeSD-Ago-1-2M2 | To express the SiAgo system, with SiAga2 carrying K12A-K13A mutations |
| pSeSD-Ago-1-2M3 | To express the SiAgo system, with SiAga2 carrying R7A-R8A-K12A-K13A mutations |
| pSeSD-AgoM1-1-2 | To express the SiAgo system, with SiAgo carrying a K142A mutation |
| pSeSD-AgoM2-1-2 | To express the SiAgo system, with SiAgo carrying a K183A mutation |

Table S2. *S. islandicus* strains constructed in the study

| Strains | Description |
|---|---|
| E:: pSeSD | *S. islandicus* E233S1 carrying the empty vector pSeSD,i.e., the CK strain |
| E:: pSeSD-Ago | E233S1 carrying the plasmid pSeSD-Ago |
| E:: pSeSD-Aga1 | E233S1 carrying the plasmid pSeSD-Aga1 |
| E:: pSeSD-Aga2 | E233S1 carrying the plasmid pSeSD-Aga2 |
| E:: pSeSD-Aga1-Aga2 | E233S1 carrying the plasmid pSeSD-Aga1-Aga2 |
| E:: pSeSD-Ago-Aga1 | E233S1 carrying the plasmid pSeSD-Ago-Aga1 |
| E:: pSeSD-Ago-Aga2 | E233S1 carrying the plasmid pSeSD-Ago-Aga2 |
| E:: pSeSD-Ago-1-2 | E233S1 carrying the plasmid pSeSD-Ago-1-2 |
| E:: pSeSD-Ago-1-2M1 | E233S1 carrying the plasmid pSeSD-Ago-1-2M1 |
| E:: pSeSD-Ago-1-2M2 | E233S1 carrying the plasmid pSeSD-Ago-1-2M2 |
| E:: pSeSD-Ago-1-2M3 | E233S1 carrying the plasmid pSeSD-Ago-1-2M3 |
| E:: pSeSD-AgoM1-1-2 | E233S1 carrying the plasmid pSeSD-AgoM1-1-2 |
| E:: pSeSD-AgoM2-1-2 | E233S1 carrying the plasmid pSeSD-AgoM2-1-2 |

Table S3. Synthesized nucleic acid in the study

| Name | Sequence |
|---|---|
| **RNA** | |
| 5P-RNA (3’-FAM) | UCAAAGCUUAGAUACCCUGGA |
| 5OH-RNA (3’-FAM) | UCAAAGCUUAGAUACCCUGGA |
| TRNA (5’-FAM) | CCUCCAGGGUAUCUAAGCUUUGAA |
| NTRNA (5’-FAM) | GUGACAGCAUCUCAUACUAGACAG |
| 5P-RNA | UCAAAGCUUAGAUACCCUGGA |
| TRNA | CCUCCAGGGUAUCUAAGCUUUGAA |
| NTRNA | GUGACAGCAUCUCAUACUAGACAG |
| **DNA** | |
| 5P-DNA (3’-FAM) | TCAAAGCTTAGATACCCTGGA |
| 5OH-DNA (3’-FAM) | TCAAAGCTTAGATACCCTGGA |
| TDNA (5’-FAM) | CCTCCAGGGTATCTAAGCTTTGAA |
| NTDNA (5’-FAM) | GTGACAGCATCTCATACTAGACAG |
| TDNA | CCTCCAGGGTATCTAAGCTTTGAA |
| NTDNA | GTGACAGCATCTCATACTAGACAG |
| Synthesized coding sequence of SiAga2 | ATGCTGAGCTCAATTACCCGTCGTTGGCGTATTAAAAAGGGCGAAGAAAGCCATGATCTGCG<br>TCTGACCTTTGTTTCATCCGGTGAATATTATGAAAAACTGCGTTCAATTGGTCTGGAACTGTT<br>TAATAAATATATCAACAGCCTGGATCTGCAGAATGAAGTGGTTCTGGCATGGGGTGCCCTGT<br>TTCATCTGGTTAGCGGTGAACTGGCCGTGAATAGCGAAAAACGTGAAGAAATTAAAAGCGAA<br>ATCCGTAAACGTCTGGAAAGCCTGATCTATCAGCTGTCAGAAAGCATGAAATCTAATTGGGA<br>ACTGGGTTTTGCAGCAAGTGTGTATCTGTATGTGCTGAAAATGATCGGTAGCAATGTTGATG<br>ATCTGGAAGATAAACTGCATACTATTCTGAAAAGTGTTAACTTTTATCGCATCGTGGGTGTTA<br>ATAAAATTCCGGAATTTCTGGGTTTTACCTTTGCGATGTTTAATGAAACCTGTCCGGATGATG<br>CATGCAAACTGTATTATTTTATGTATAAAAAGATCAACAACGAAGATATCAACTTTGAACAGAA<br>ACAGTTTAACAAAAACGATCTGATTGAATCCCTGCTGGCATATTATATTTATGGTGAAACCAG<br>CGGCCTGCAGCTGAAACAGCTGTGGATTCAAATTATTCGTGAAGTTATGGAAAAAGATGTGA<br>ACGAATATATGAAAAACTATGAACCTTATGGCGGTGAATATTATAGTAATGAAGTTTTTGTGC<br>CGAAAGAACGTCTGTTTCTGCTGCTGATTCTGATGAAACTGCTGAATCTGGATAAAGTGGTG<br>TATGTGATTGGTCTGTCTAATGAAAAAGATATGAAAGAAATCCTGGAACAGGATAAAGAAATT<br>AAAAAGGCAAACCGTCGTGCAACTGTGATCCTGGTTTTTAGCCTGATCAGTATTACCCTGCT<br>GATTATTTGGGGTCTGGTTATTCCGTTTATGTTTCCTAGCCTGTTTTTTACCCTGCTGCATAGT<br>ATTGTTGCTATTGTTGTTGTTATCATCTCTTATATCCTGGGCCTGATTAGTATTATTATTGAAAT<br>TCTGAAAAAGCTGCTGGATTGGGTTCGCGTTAAAAAGGTTAATTAA |

Table S4. Primers used in the study

| Name | Sequence | Description |
|---|---|---|
| Ago-F_Nhe1 | CTAGCTAGCATGAGTGAGTATGCCACTATATTAC | To insert the *ago* gene into pSeSD and pET30aN |
| Ago-R_Not1 | AAATATGCGGCCGCTTAAAGTACATCCCGCAC | |
| AgoM1-F | CATGACAAAACAGCAGCTGCATTGATAGGTAG | Overlapping primers to generate the K142A mutation |
| AgoM1-R | AATGCAGCTGCTGTTTTGTCATGAAATTCCTTAG | |
| AgoM2-F | GCTGCAGGAGGTACTCCATGGATAG | Overlapping primers to generate the K183A mutation |
| AgoM2-R | AGTACCTCCTGCAGCTGCAATTAATTGCACTGCAAG | |
| Aras-F_Sma1 | ATGCCCCGGGATGTTAAACAAGTTAGG | To amplify the Ago expression cassette |
| Aga1-F_Nde1 | GAATGAGGTGAAGCTCATATGGTTTTAGAATCTAATATG | To insert *aga1* into pSeSD |
| Aga1-R_Not1 | GGTGATGATGATGTGCGGCCGCTTAAAAGGGAGTAGGCAACTTTTC | |
| Aga1-F_Nhe1 | CTAGCTAGCATGGTTTTAGAATCTAATATG | To insert *aga1* into pET30aN |
| Aga1-R_Not1-2 | AAATATGCGGCCGCTTAAAAGGGAGTAGGCAACTTTTC | |
| Aga1-R_Xho1 | CCGCTCGAGTTAAAAGGGAGTAGGCAACTTTTC | To insert *aga1* into pET21d |
| Aga2-F_Nde1 | GAGAATGAGGTGAAGCTCATATGCTTTCTAGTATTACTCG | To insert the native *aga2* gene into pSeSD |
| Aga2-R_NotI | GTGGTGATGATGATGTGCGGCCGCTTAATTCACCTTTTTTACACG | |
| Aga2-F_Nhe1 | CTAGCTAGCATGCTTTCTAGTATTACTCG | To insert the native *aga2* gene into pET30aN |
| Aga2-R_Not1-2 | AAATATGCGGCCGCCTAATTCACCTTTTTTACACGC | |
| O-Aga2-F_Nde1 | GGAATTCCATATGCTGAGCTCAATTACCC | To insert the optimized *aga2* gene into pMAL-c5x |
| O-Aga2-R_Not1 | ATAAGAATGCGGCCGCTTAATTACCTGCAGGGAATTC | |
| MBP-F_BamH1 | CGGGATCCATGAAAATCGAAGAAGGTAAAC | To amplify the *mbp-aga2* fusion gene |
| MBP(HF)-F_Nde1 | GGAATTCCATATGAAAATCGAAGAAGGTAAAC | To amplify the *mbp-aga2* fusion gene for expressing His-tag free protein |
| O-Aga2(HF)-R_Xho1 | CCGCTCGAGTTAATTACCTGCAGGGAATTC | |
| MBP(HF)-R_Xho1 | CCGCTCGAGTTAAGTCTGCGCGTCTTTC | To amplify the *mbp* gene |
| O-Aga2ΔC-R_Xho1 | CCGCTCGAGTCACACAGTTGCACGACG | To amplify the truncated *aga2* gene |
| Aga2M1-F | ACTGCAGCATGGAGGATTAAAAAAGG | Overlapping primers to generate the R7A-R8A mutations |
| Aga2M1-R | CCTCCATGCTGCAGTAATACTAGAAAGC | |
| Aga2M2-F | GATTGCAGCAGGAGAAGAAAGCCAC | Overlapping primers to generate the K12A-K13A mutations |
| Aga2M2-R | TCTCCTGCTGCAATCCTCCATCTTCG | |

| | | |
|---|---|---|
| Aga2M3-R | CTTCTCCTGCTGCAATCCTCCATGCTGCAGTAATAC | Overlapping primers to generate the R7A-R8A-K12A-K13A mutations |
| O-Aga2ΔCM1-F | CCGCTGCTTGGCGTATTAAAAAGGGC | Overlapping primers to generate the R7A-R8A mutations |
| O-Aga2ΔCM1-R | TACGCCAAGCAGCGGTAATTGAGCTCAGCAT | |
| O-Aga2ΔCM2-F | GCAGCGGGCGAAGAAAGCCATGATC | Overlapping primers to generate the K12A-K13A mutations |
| O-Aga2ΔCM2-R | CTTTCTTCGCCCGCTGCAATACGCCAACGACGGG | |
| O-Aga2ΔCM3-R | CTTTCTTCGCCCGCTGCAATACGCCAAGCAGCGGTAATTG | Overlapping primers to generate the R7A-R8A-K12A-K13A mutations |
